# Investigating macroecological patterns in coarse-grained microbial communities using the stochastic logistic model of growth

**DOI:** 10.1101/2023.03.02.530804

**Authors:** William R. Shoemaker, Jacopo Grilli

## Abstract

The structure and diversity of microbial communities is intrinsically hierarchical due to the shared evolutionary history of their constituents. This history is typically captured through taxonomic assignment and phylogenetic reconstruction, sources of information that are frequently used to group microbes into higher levels of organization in experimental and natural communities. Connecting community diversity to the joint ecological dynamics of the abundances of these groups is a central problem of community ecology. However, how microbial diversity depends on the scale of observation at which groups are defined has never been systematically examined. Here, we used a macroecological approach to quantitatively characterize the structure and diversity of microbial communities among disparate environments across taxonomic and phylogenetic scales. We found that measures of biodiversity at a given scale can be consistently predicted using a minimal model of ecology, the Stochastic Logistic Model of growth (SLM). This result suggests that the SLM is a more appropriate null-model for microbial biodiversity than alternatives such as the Unified Neutral Theory of Biodiversity. Extending these within-scale results, we examined the relationship between measures of biodiversity calculated at different scales (e.g., genus vs. family), an empirical pattern predicted by the Diversity Begets Diversity (DBD) hypothesis. We found that the relationship between richness estimates at different scales can be quantitatively predicted assuming independence among community members.Contrastingly, only by including correlations between the abundances of community members (e.g., as the consequence of interactions) can we predict the relationship between estimates of diversity at different scales. The results of this study characterize novel microbial patterns across scales of organization and establish a sharp demarcation between recently proposed macroecological patterns that are not and are affected by ecological interactions.

## Introduction

An essential feature of microbial communities is their heterogeneous composition. A single environmental sample typically has a high richness, harboring hundreds to thousands of community members [1–3]. This high level of richness reaches an astronomical quantity at the global level, as scaling relationships and models of biodiversity predict upwards of one trillion (∼10^12^) species on Earth [4, 5]. Even experimental communities in laboratory settings with a single carbon source can harbor ≥40 community members, culminating in a total richness numbering in the hundreds among replicate communities (e.g., [6]). This richness contributes to the sheer diversity of microbial communities, a challenge for researchers attempting to identify the general principles that govern their dynamics and composition.

While richness estimates of microbial communities are undoubtedly high, the choice of assigning a community member to a given taxon remains intrinsically arbitrary. This arbitrariness remains regardless of whether the definition of a taxon is based on physiological attributes measured in the laboratory, entire genomes (i.e., metagenomics), or single-gene amplicon-based methods (i.e., 16S rRNA annotation). Despite their methodological differences, these approaches can all be viewed as different ways to cluster individuals within a community into groups. To contend with the sheer richness of microbial communities, researchers frequently rely on annotation-based approaches, i.e., by summing the abundances of community members that belong to the same group at a given taxonomic scale (e.g., genus, family, etc.). This approach pares down communities to a size that is amenable for the visualization of individual groups and allows for questions of scale-dependent community reproducibility to be addressed [6–13].

This movement towards performing analyses of diversity at various taxonomic scales raises the question of how the composition of a community at one scale relates to that at another. To address these questions, researchers have examined the relationship between biodiversity measures at different scales in order to pare down the set of ecological mechanisms that plausibly govern community composition. Specifically, recent efforts have found that microbial richness/diversity within a given taxonomic group (e.g., genus) is typically positively correlated with the richness/diversity among the remaining groups (e.g., family) [9, 14], an empirical pattern that aligns with the predictions of the Diversity Begets Diversity hypothesis (DBD) [15–17]. Evidence of the DBD hypothesis has historically been attributed to the construction of novel niches within a community through member interactions [15, 18], with similar mechanisms having been proposed to explain the existence of a positive relationship in microbial communities [14]. However, we still lack a quantitative understanding of how community composition at one scale should relate to that of another. Proceeding towards this goal requires two elements: 1) a systematic approach to grouping community members and 2) an appropriate null model for the composition of communities.

The operation of grouping the components of a system into a smaller number (e.g., merging read counts of OTUs to the family level in a community) is known in the physical sciences as coarse-graining. This formalism defines our systematic approach to grouping community members. While it is often not explicitly acknowledged as such, coarse-graining is a core concept in the microbial life sciences [19]. By smoothing over microscopic details at a lower level of biological organization in order to make progress at a higher level, the concept of coarse-graining has contributed towards the development of effective models of physiological growth [20, 21], evolutionary dynamics [22, 23], and the dependence of ecosystem properties on the diversity of underlying communities [24]. Coarse-graining has even been used to glean insight into the question of whether “species” as a unit has meaning for microorganisms, as modeling efforts have found that the operation permits the delimitation of distinct taxonomic groups when the resource preferences of community members are structured [25]. These theoretical and empirical efforts suggest that coarse-graining may provide an appropriate framework for investigating patterns of diversity and abundance within and between taxonomic scales of observation.

When evaluating the novelty of an empirical pattern it is useful to identify an appropriate null model for comparison [26–28]. Prior research efforts have demonstrated the novelty of the fine vs. coarse-grained relationship by contrasting inferences from empirical data with predictions obtained from the Unified Neutral Theory of Biodiversity (UNTB) [14, 29–33]. These predictions generally failed to reproduce slopes inferred from empirical data [14], implying that the fine vs. coarse-grained relationship represents a novel macroecological pattern that cannot be quantitatively explained by existing null models of ecology. However, the task of identifying an appropriate null model for comparison is not straightforward. Rather, the question of what constitutes an appropriate null model remains a persistent topic of discussion in community ecology [26, 34–37]. Here we take the view that a null model is appropriate for examining the relationship between two observables (e.g., community diversity at different scales) if it was capable of quantitatively predicting each observable (e.g., community diversity at one scale). By this standard, the UNTB is an unsuitable choice as a null as it generally fails to capture basic patterns of microbial diversity and abundance at any scale [38–40]. One relevant example is that the UNTB predicts that the distribution of mean abundances of community members across sites is extremely narrow (i.e., converging to a delta distribution as the number of sites increases), whereas empirical data tends to follow a broad lognormal distribution [40]. Contrastingly, recent efforts have determined that the predictions of a model of self-limiting growth with environmental noise, the Stochastic Logistic Model (SLM), is capable of quantitatively capturing multiple empirical macroecological patterns in observational and experimental microbial communities [40–46]. The stationary solution of this model predicts that the abundance of a given community member across sites follows a gamma distribution [40], a result that provides the foundation necessary to predict macroecological patterns among and between different taxonomic and phylogenetic scales.

In this study, we evaluated macroecological patterns of microbial communities across scales of evolutionary resolution. To limit potential biases that may result due to taxonomic annotation errors and to use all available data, we investigated the macroecological consequences of coarse-graining by developing a procedure that groups community members using the underlying phylogeny in addition to relying on taxonomic assignment. We used data from the Earth Microbiome Project (EMP), a public catalogue of microbial community barcode data, to ensure the generality of our findings and their commensurability with past research efforts. First, we assessed the extent that microbial diversity varies as the abundances of community members are coarse-grained by phylogenetic distance and taxonomic rank. The results of these analyses lead us to consider whether the predictive capacity of the gamma distribution remained robust under coarse-graining, a prediction that we quantitatively evaluated among community members and then extended to predict overall community richness and diversity. The accuracy of the gamma distribution provided the necessary motivation to test whether the gamma distribution was capable of predicting the relationship between fine and coarse-grained estimates, the empirical pattern that has been interpreted as evidence for the DBD hypothesis. Together, these analyses present evidence of the scale invariance of macroecological patterns in microbial communities as well as the applicability of the gamma distribution, the stationary distribution of the SLM, as a null model for evaluating the novelty of macroecological patterns of microbial biodiversity.

## Results

### The macroecological consequences of phylogenetic and taxonomic coarse-graining

While microbial communities are often coarse-grained into higher taxonomic scales, their effect on measures of biodiversity and the underlying phylogeny are rarely examined. Before proceeding with the full analysis using public 16S rRNA amplicon data from the EMP, we elected to quantify the fraction of remaining community members across coarse-graining thresholds, a reflection of the extent that coarse-graining reduces global richness and the relation between taxonomic and phylogenetic coarse-graining. We first defined a coarse-grained group *g* as the set of OTUs that have the same assigned label in a given taxonomic rank out of *G* groups (e.g., *Pseudomonas* at the genus level) or are collapsed when the phylogeny is truncated by a given root-to-tip distance (Figs. 1, S1). The relative abundance of group *g* in site *j* is defined as *x*_*gj*_ =Σ _*i*∈*g*_ *x* _*i,j*_.

**Figure 1.**
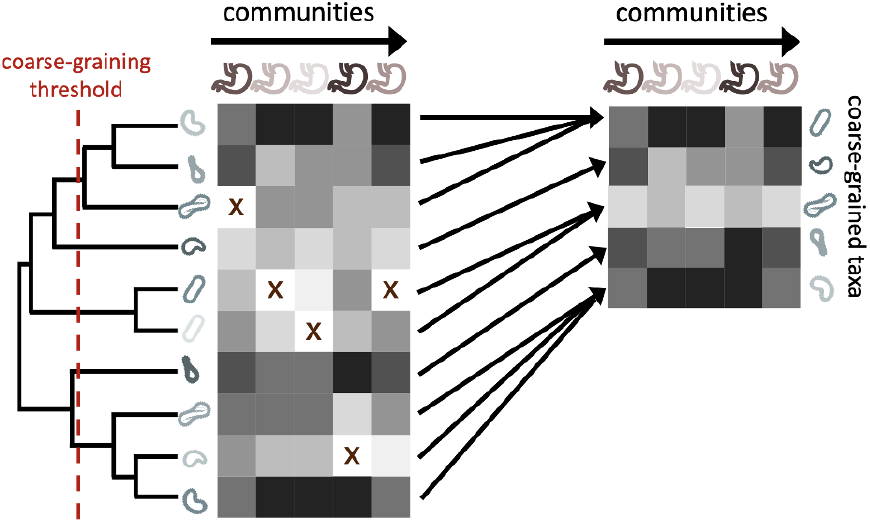
The process of coarse-graining abundances using the phylogeny. Taxonomic assignment in 16S rRNA amplicon sequence data provided the opportunity to investigate how properties of communities vary at different taxonomic scales. The most straightforward means of coarse-graining here is to sum the abundances of OTUs/ASVs that belong to the same taxonomic group. Amplicon data-based studies provide information about the shared evolutionary history of its constituents, information that can be leveraged by the construction of phylogenetic trees. A coarse-graining procedure can be defined that is analogous to one based on taxonomy, where a phylogenetic distance is chosen, and terminal nodes are collapsed if their distance to a common ancestor is less than the prescribed distance.

We found that coarse-graining had a drastic effect on the total number of community members within an environment, reducing global richness by ∼90% even at just the genus level (Fig. S2a). By coarse-graining over a range of phylogenetic distances, we found that a fraction of coarse-grained community members comparable to that of genus-level coarse-graining occurred at a phylogenetic distance of ∼0.1 (Fig. S2b). This distance translated to only ∼3% of the total distance of the tree, meaning that the majority of OTUs were coarse-grained over a minority of the tree. This pattern was likely driven by the underlying structure of microbial phylogenetic trees, where most community members have short branch lengths [47]. This result suggests that while coarse-graining communities to the genus or family level substantially reduces global richness, it does so without coarse-graining the majority of the evolutionary history captured by the phylogeny. Assuming that phylogenies capture ecological changes that occur over evolutionary time, this detail implies that ecological divergence that is captured by the phylogeny should be retained even when communities are considerably coarse-grained.

With our coarse-graining procedures established, we proceeded with our macroecological investigation. Recent efforts have found that the distribution of abundances of a given ASV/OTU maintained a consistent statistically similar form across independent sites and time, a pattern known as the Abundance Fluctuation Distribution [40–42, 48, 49]. By coarse-graining empirical AFDs and rescaling them by their mean and variance across sites (i.e., standard score), we found that AFDs from the human gut microbiome retained their shape across phylogenetic scales (Fig. 2a). This pattern of invariance held across environments for both phylogenetic and taxonomic coarse-graining (Figs. S3, S4), suggesting that empirical AFDs can likely be described by a single probability distribution.

**Figure 2.**
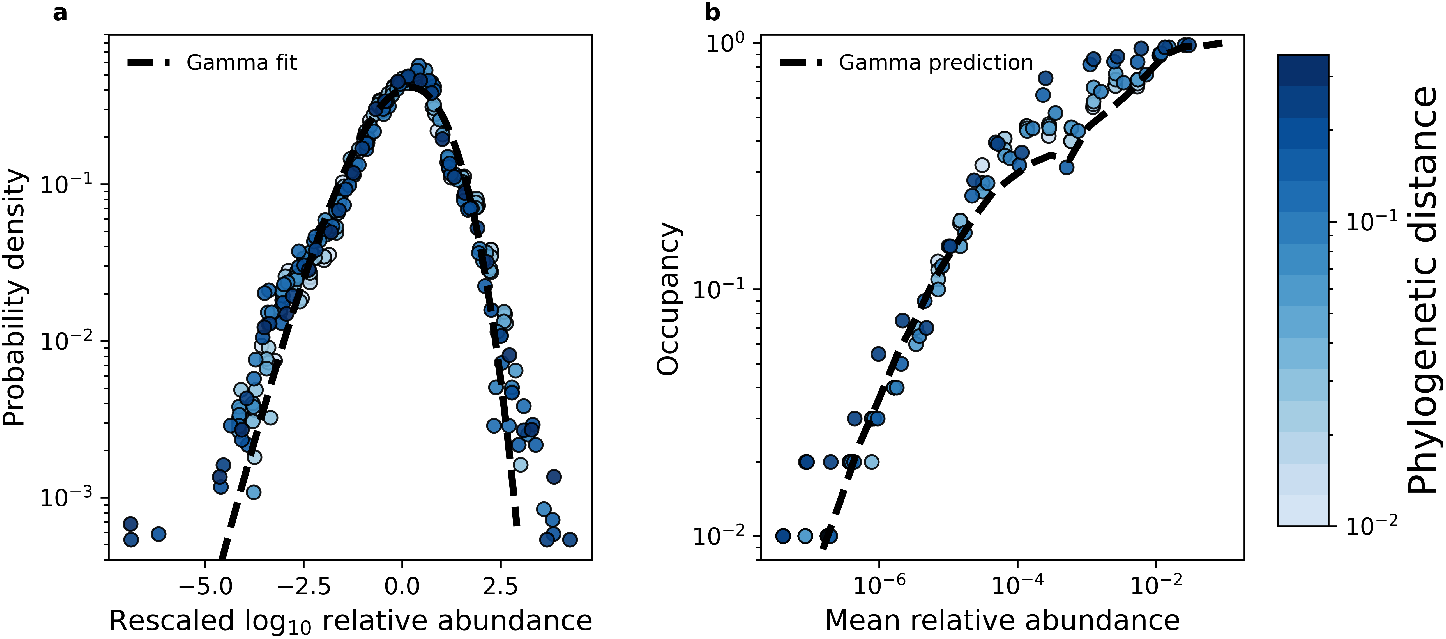
The shape of the AFD remained qualitatively invariant under coarse-graining. **a)** Under phylogenetic coarse-graining the general shape of the AFD for OTUs that were present in all sites (i.e., an occupancy of one) remained qualitatively invariant. **b)** Similarly, the shape of the relationship between the mean coarse-grained abundance across hosts and occupancy across sites did not tend to vary. Predictions obtained from the gamma distribution are capable of capturing the relationship between the mean abundance and occupancy, suggesting that the gamma distribution remains a useful quantitative null model under coarse-graining. All data in this plot is from the human gut microbiome.

It has been previously demonstrated that empirical microbial AFDs are well-described by a gamma distribution that is parameterized by the mean relative abundance 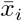 and the shape parameter 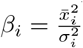 (equal to the squared inverse of the coefficient of variation [40]). This distribution can be viewed as the stationary distribution of a Stochastic Logistic Model (SLM) of growth, a mathematical model that successfully captures macroecological patterns of microbial communities across both sites and time [40, 41, 44] (Eq. 5 in Materials and Methods).

Using this result, we determined whether the gamma distribution sufficiently characterized coarse-grained AFDs. In order to accomplish this task, it is worth noting that we do not directly observe *x*_*i*_. Rather, our ability to observe a community member is dependent on sampling effort (i.e., total number of reads for a given site). To account for sampling, one can derive a form of the gamma distribution that explicitly accounts for the sampling process, obtaining the probability of obtaining *n* reads out of *N* total reads belonging to a community member 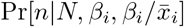 (Materials and Methods, [40]). Given that *n* = 0 for a community member we do not observe, we defined the fraction of *M* sites where a community member was observed (i.e., occupancy, *o*_*i*_) as

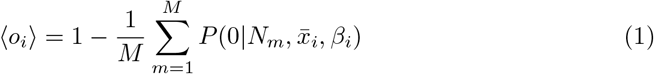

We then compared this prediction to observed estimates of occupancy to assess the accuracy of the gamma distribution across coarse-grained thresholds. We found that Eq. 1 generally succeeded in predicting observed occupancy across phylogenetic and taxonomic scales for all environments (Figs. S5, S6). We then determined whether the gamma distribution was capable of predicting the relationship between macroecological quantities. One such relationship is that the occupancy of a community member should increase with its mean abundance, known as the abundance-occupancy relationship [50]. This pattern has been found across microbial systems [51–53] and can be quantitatively predicted using the gamma distribution [40]. We see that this relationship is broadly captured across taxonomic and phylogenetic scales for all environments (Figs. 2b, S7 S8). This result implies that the ability to observe a given taxonomic group was primarily determined by its mean abundance across sites and the sampling effort within a site, regardless of one’s scale of observation. In contrast, under the assumption of demographic indistinguishability under the UNTB we would expect the mean abundance distribution to be extremely narrow, following a delta distribution. Under the SLM, the variation in mean relative abundances we observed implies that the carrying capacities of community members vary over multiple orders of magnitude. We also note that at high mean abundances our predictions show slight variation, which is likely driven by variation in the shape parameter *β* (Figs. 2b). In contrast with these results, the gamma distribution was unable to predict the variance of occupancy under both taxonomic and phylogenetic coarse-graining (Eq. 11; Figs. S9, S10), the implications of which we will address in a later section.

To quantitatively assess the accuracy of the gamma distribution we calculated the relative error of our mean occupancy predictions (Eq. 12) for all coarse-graining thresholds. We found that the mean logarithm of the error only slightly increased for the initial taxonomic and phylogenetic scales, where it then exhibited a sharp decrease across environments (Fig. S11a,b). The error then only began to decrease once the community became highly coarse-grained, harboring a global richness (union of all community members in all sites for a given environment) *<* 20. This result means that, if anything, the accuracy of the gamma distribution only improved with coarse-graining.

### Reconciling coarse-graining and the predictions of the gamma distribution

The consistent predictive success of the gamma distribution under coarse-graining raises the question of why it remains a sufficient null model. The sum of independent gamma distributed random variables only returns a gamma through analytic calculation if all random variables have identical rate parameters 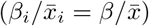, a requirement that microbial communities clearly do not meet since they typically harbor broad mean abundance distributions. Given that a gamma AFD cannot predict the distribution of correlations between AFDs [40], it is first worth examining whether the degree of dependence between AFDs shapes coarse-grained variables. We first consider the relation between the variance of the sum and the sum of variances.

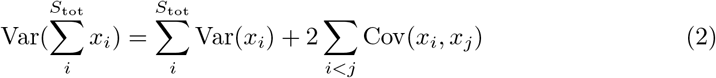

By plotting 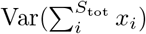 against 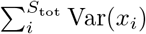 across coarse-grained thresholds, we found that the contribution of covariance to individual coarse-grained taxa was weak, suggesting that the statistical moments at higher scales can be approximated by those at lower scales (Figs. S12, S13). Similar conclusions can be drawn by plotting the variances as a ratio, with slight deviations above a ratio of one, suggesting that coarse-grained variance was slightly higher (Figs. S14, S15). These results are consistent with previous efforts demonstrating that the strongest correlations between AFDs are typically concentrated among pairs of closely related community members (i.e., low phylogenetic distance) [54], implying that the effects of correlation should dissipate when communities are coarse-grained. Given that the variance of the sum can be approximated by the sum of the variances and that, by definition, the mean of a sum is the sum of the means, it is reasonable to propose that the statistical moments of coarse-grained AFDs are sufficient to characterize the distribution.

Finally, while we know of no general closed-form solution for the sum of independent gamma distributed random variables with different rate parameters (equivalent to considering the convolution of many AFDs with different carrying capacities), progress has been made towards obtaining suitable approximations [55–59]. This body of work includes an analysis demonstrating that a single gamma distribution can provide a suitable approximation to the distribution of the sum of many gamma random variables with different rate parameters [60]. In summary, the gamma distribution appears to successfully captures patterns of biodiversity under taxonomic and phylogenetic coarse-graining because the sum of multiple gamma distributions can be approximated by a single gamma distribution.

### Predicting measures of richness and diversity within a coarse-grained scale

Given that the presence or absence of a community member is used to estimate community richness, a measure previously used to make claims about patterns of microbial diversity across taxonomic scales [14], we can visualize the sufficiency of the gamma distribution by predicting the mean richness within an environment at a given coarse-grained scale (Eq. 13). Likewise, we can use the entirety of the distribution of read counts to predict the diversity within a site, a measure that reflects richness as well as the distribution of abundances within a community (Eq. 14), analytic predictions that we validated through simulations (Fig. S16). We note that we observe consistent deviations between the analytic predictions of the variance of diversity and simulation results. These deviations are likely driven by small deviations in predictions of the second moment of diversity, which are slight for individual community members, but become considerable when terms are summed over hundreds or thousands of community members.

Focusing on the human gut microbiome as an example, we found that we can predict the typical richness of a community across phylogenetic scales using the gamma distribution (Fig. 3a). Similar results were obtained when we repeated the our analysis for predicted diversity (Fig. 3b). By examining all nine environments we found that despite the dissimilarity in environments, we were able to predict mean richness and diversity in the face of coarse-graining (Fig. 3c,d). In contrast, the UNTB failed to predict richness (Fig. S18). The results of this analysis suggest that the composition of microbial communities remained largely invariant under coarse-graining and that the gamma distribution remained a suitable null model for predicting mean community measures across coarse-grained scales. Identical results were obtained for taxonomic coarse-graining (Fig. S17).

**Figure 3.**
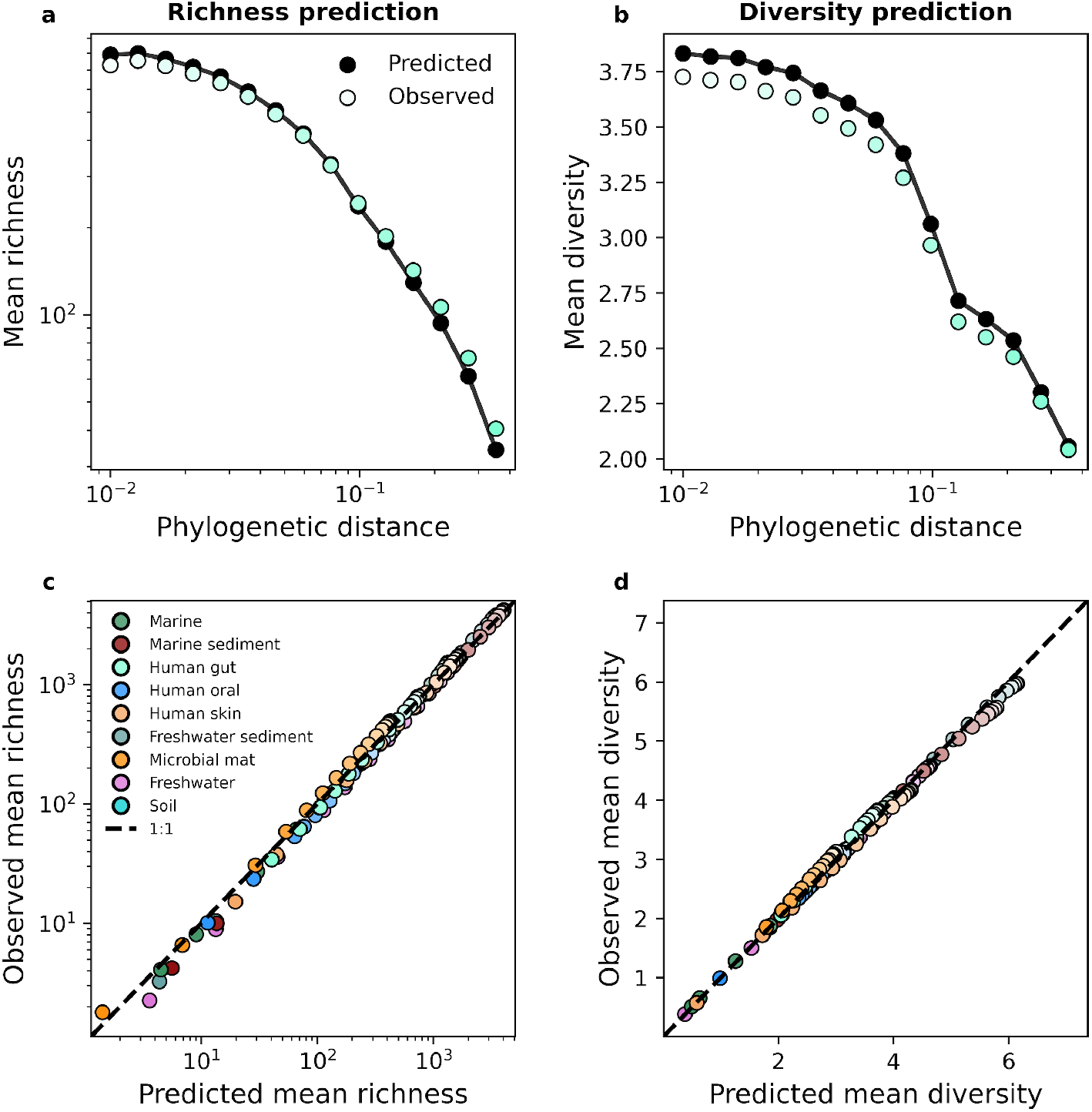
The gamma distribution successfully predicted mean richness and diversity under phylogenetic coarse-graining. **a)** The expected richness derived from the gamma distribution (Eq. 13) was capable of predicting richness across phylogenetic coarse-graining scales, as illustrated by data from the human gut. **b)** Predictions remained successful across all environments, suggesting that a minimal model of zero interactions was sufficient to predict observed properties of community composition **c**,**d)** Similarly, predictions of expected diversity (14) also succeeded across coarse-graining scales for all environments. The shade of a color of a given datapoint represents the phylogenetic distance used for coarse-graining, with lighter colors representing finer scales and darker colors representing coarser scales.

Turning to higher-order moments, we examined the variance of richness and diversity across sites. Using a similar approach that was applied to the mean, we derived analytic predictions for the variance (Eq. 17). With the human gut as an example, we see that analytic predictions typically fail to capture estimates of variance obtained from empirical data for phylogenetic coarse-graining (Fig. 4a,b). This lack of predictive success was consistent across environments (Fig. 4c,d), implying that a model of independent community members with gamma distributed abundances was insufficient to capture the variance of measures of biodiversity. A major assumption made in our derivation was that community members are independent, an assumption that is unjustified given that the gamma distribution has been previously shown to be unable to capture the empirical distribution of correlations in the AFDs of community members [40]. To attempt to remedy this failed prediction, we again turned to the law of total variance by estimating the covariance of richness and diversity from empirical data and adding the covariance to the predicted variance for each measure. We found that the addition of this empirical estimate was sufficient to predict the observed variance in the human gut (Fig. 4a,b) as well as across environments (Fig. 4e,f), implying that the underlying model is fundamentally correct for predicting the first moment of measures of biodiversity but cannot capture the correlations necessary to explain higher statistical moments such as the variance. Identical results were again obtained with taxonomic coarse-graining (Fig. S19).

**Figure 4.**
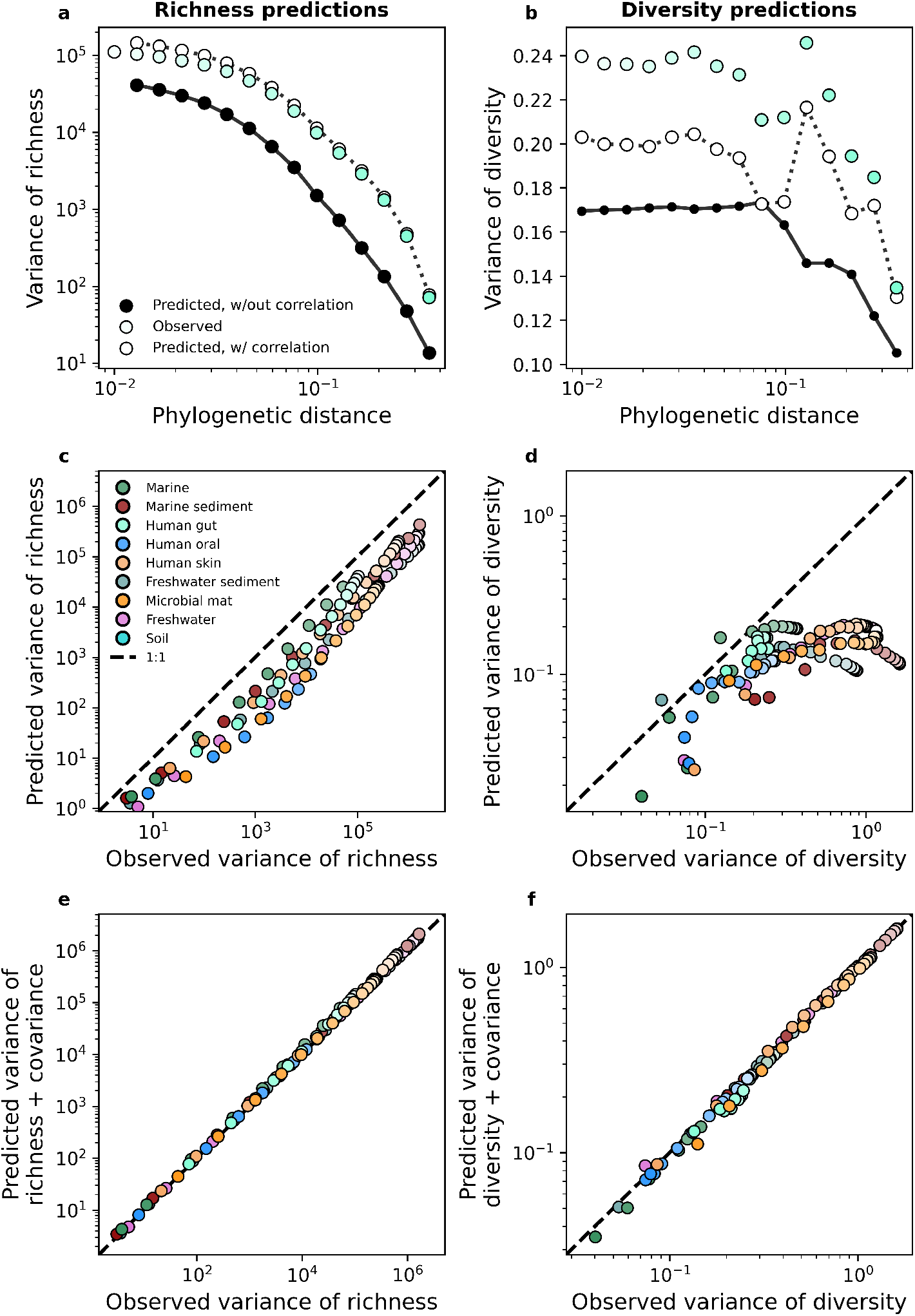
The gamma distribution only predicts the variance of richness and diversity under phylogenetic coarse-graining when covariance is included. **a,b)** In contrast with the mean, the variance of richness and diversity estimates predicted by the gamma distribution (Eq. 17) failed to capture empirical estimates from the human gut. Predictions are only comparable when empirical estimates of covariance are included in the predictions of the gamma distribution, meaning that dependence among community members is essential to describe the variation in measures of biodiversity across communities **c**,**d)** This lack of predictive success was constant across environments, **e**,**f)** though the addition of covariance consistently improves our analytic predictions. The colorscale used here is identical to the colorscale used in Fig. 3.

### Predicting patterns of richness and diversity between fine and coarse-grained scales

Our predictions of the statistical moments of richness and diversity using the gamma distribution provided the foundation necessary to investigate macroecological patterns between different taxonomic and phylogenetic scales. One such prominent pattern is the relationship between the fine-grain richness/diversity within a given coarse-grained group vs. the coarse-grained richness/diversity among all remaining groups (e.g., the number of classes within Firmicutes vs. the number of phyla excluding the phylum Firmicutes), a pattern that has been purported to demonstrate the existence of DBD processes in microbial systems. Before continuing, we note that the acronym DBD technically refers to the hypothesis that such positive relationships reflects the existence of ecological interactions through which coarse-grained diversity bolsters the accumulation of fine-grained diversity (e.g., niche construction [16, 61]). Since we are primarily interested in the predictive power of an empirically-validated null model of biodiversity, we distinguish between DBD as a hypothesis and DBD as an empirical pattern by referring to the slope as the fine vs. coarse-grained relationship throughout the remainder of this manuscript.

The fine vs. coarse-grained relationship can be quantified as the slope of the relationship between the fine-grained richness within a given coarse-grained group *g* (*S* _*g,m*_) and the richness in the remaining *G*−1 coarse-grained groups: *S*_*g,m*_ ∝*αS*_*G/g,m*_, where *G*\*g* denotes the exclusion of group *g* and *α* is the slope of the relationship. This formulation was proposed in Madi et al. and to ensure commensurability we adopted it here [14]. Furthermore, keeping with the approach used by Madi et al., fine and coarse-grained measures were compared across increasing taxonomic and phylogenetic scales (e.g., OTU vs. genus, genus vs. family, etc.), [14]. Using Eq. 1, we then defined each of these estimators in terms of the sampling form of the gamma distribution while accounting for sampling

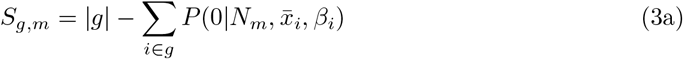

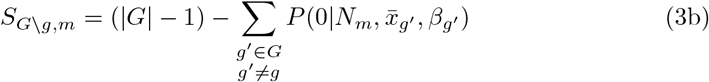

Similarly, we used Eq. 14 to derive predictions for fine and coarse-grained diversity.

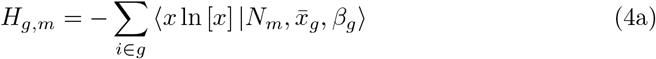

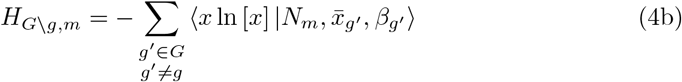

By repeating this calculation for all *M* sites, we obtained vectors of coarse and fine-grained richness estimates for group *g* from which we inferred the slope of the fine vs. coarse-grained relationship through ordinary least squares regression. By repeating this process for all *G* groups we obtained a distribution of slopes that can be directly compared to those obtained from empirical data. We include a conceptual diagram visualizing this process in the supplement S20.

Before performing a direct comparison, we first note the features of the empirical slopes and how they pertain to the predictions we obtained. By examining the distribution of empirical slopes pooled over all coarse-graining thresholds for each environment, we found that they were rarely less than zero (Figs. 5a, S21a). The few negative slopes inferred from empirical data were extremely small, having absolute values *<* 10^*-*4^ and could be treated as zeros. Furthermore, the distribution of slopes follows the same form across environments, suggesting that the slope of the fine vs. coarse-grained relationship reflects a general feature of community sequence data rather than the ecology of specific environments. Like the empirical slopes, the gamma distribution virtually always predicted a positive slope for all environments for both taxonomic and phylogenetic coarse-graining. This paucity of negative slopes suggests that the prediction of the alternative to the DBD hypothesis, the Ecological Controls hypothesis [62], is virtually absent in empirical data and cannot be generated from an empirically validated null model of microbial biodiversity.

**Figure 5.**
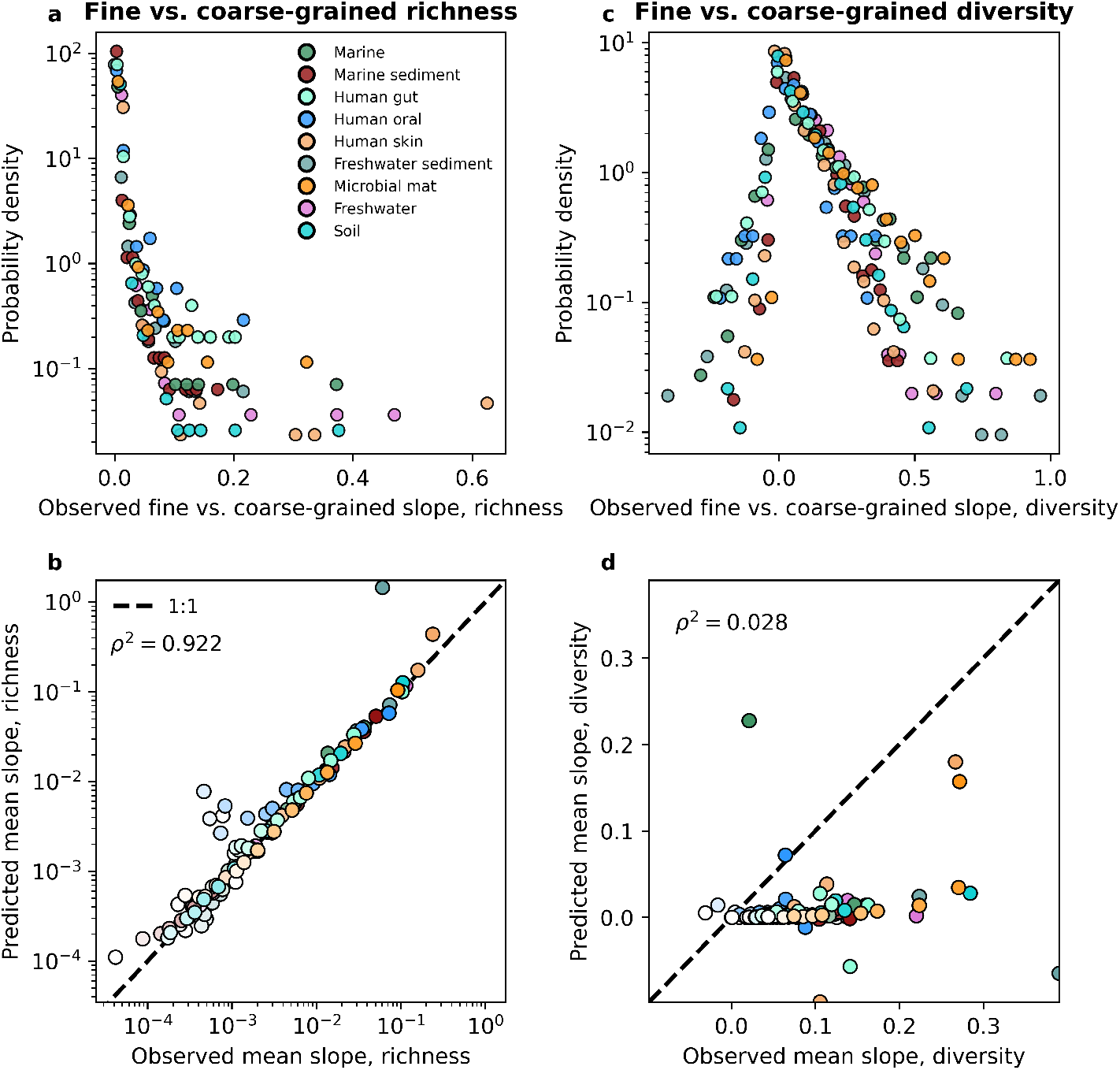
The slope of the fine vs. coarse-grained relationship for richness could be predicted by the gamma distribution, but was novel for estimates of diversity. **a,b)** The predictions of the gamma distribution (Eq. 3) successfully reproduced observed fine vs. coarse-grained richness slopes across scales of phylogenetic coarse-graining. **c**,**d)** In contrast, the predictions of the gamma distribution failed to capture diversity slopes (Eq. 4). The colorscale used here is identical to the colorscale used in Fig. 3. Squared Pearson correlation coefficients (*ρ*^2^) are computed over all slopes for all taxa across all coarse-graining scales.

However, only observing positive slopes does not necessarily provide support to the DBD hypothesis. A direct comparison of slopes predicted from the gamma distribution to those inferred from empirical data is necessary to determine whether the predictions of DBD lie outside what can be reasonably captured by an interaction-free model such as the SLM. To evaluate the novelty of the slope of the fine vs. coarse-grained relationship we compared the values of observed slopes to those obtained from the interaction-free SLM. We found that the predictions of the gamma distribution closely matched the observed slopes across environments for both taxonomic and phylogenetic coarse-graining (Figs. S22, S23). We consolidated these results by taking the mean slope for a given coarse-grained level, from which we see that the mean slope predicted by the gamma distribution does a reasonable job capturing empirical slopes across environments (Figs. 5b, S21b). These results indicate that we should expect to see a positive relationship between richness estimates at different scales and that the relationships we observe can be quantitatively captured by a gamma distributed AFD. It is worth noting that the slope of the fine vs. coarse-grained relationship could be sufficiently predicted even though the gamma distribution only succeeded at predicting mean richness, suggesting that higher order statistical moments, and by extension interactions between community members, are unnecessary to quantitatively capture the positive relationship observed between fine and coarse-grained estimates of richness.

As a point of comparison we predicted the slope of the fine vs. coarse-grained relationship for richness using a UNTB model [63] (Supporting Information). We found that generally the UNTB slopes deviated from those obtained from empirical data, exhibiting far more variation around the 1:1 line than observed for the SLM (Figs. S24, S25). By examining the mean slope we found that predictions from the UNTB tended to systematically underpredict the observed slope under both taxonomic and phylogenetic coarse-graining (Fig. S26). Directly comparing the mean relative error of the UNTB predictions to those of the SLM confirms these observations, as the UNTB predictions tended to have larger errors by an order of magnitude (Figs. S27, S28). To summarize, in contrast to the SLM, the UNTB cannot predict the slope of the fine vs. coarse-grained relationship for richness.

While richness is a widespread and versatile estimator that is commonly used in community ecology, neglects considerable information by focusing on presences and absences instead of the entirety of the distribution of abundances. To rigorously test the predictive power of the gamma distribution it was necessary to evaluate the fine vs. coarse-grained relationship for diversity. We again found that disparate environments had similar distributions of slopes from empirical data (Figs. 5c, S21c), suggesting that the slope of the relationship is likely a general property of microbial communities rather than an environment-specific pattern. However, unlike richness, diversity predictions obtained from the gamma distribution generally failed to capture observed slopes, as the squared correlation between observed and predicted slopes can be less than that of richness by over an order of magnitude (Figs. 5d, S29, S30, S21d). Here we see where the predictions of an interaction-free SLM succeeded and failed to predict observed macroecological patterns.

Given that the gamma distribution failed to predict the observed diversity slope, it is worth evaluating whether additional features could be incorporated to generate successful predictions. A notable omission is that there is an absence of interactions between community members in the SLM, meaning that we were unable to predict correlations between community member abundances. However, while considerable progress has been made (e.g., [11]), predicting the observed distribution of correlation coefficients between community members while accounting for sampling remains a non-trivial task. Given that the gamma distribution succeeded at predicting other macroecological patterns, we elected to perform a simulation where a collection of sites was modeled as an ensemble of communities with correlated gamma-distributed AFDs with the means, variances, correlations, and total depth of sampling set by estimates from empirical data (Materials and methods). By including correlations between AFDs into the simulations, the statistical outcome of ecological interactions between community members, we were able to largely capture observed fine vs. coarse-grained diversity slopes (Figs. 6, S31, S32, S33). These results suggest that rather than diversity at a fine-scale begetting diversity at a coarse-scale, the correlations that exist at a fine-scale (e.g., genus) contribute to measures of biodiversity at the nearest coarse-grained scale (e.g., family), resulting in a positive relationship between measures of diversity at different scales.

**Figure 6.**
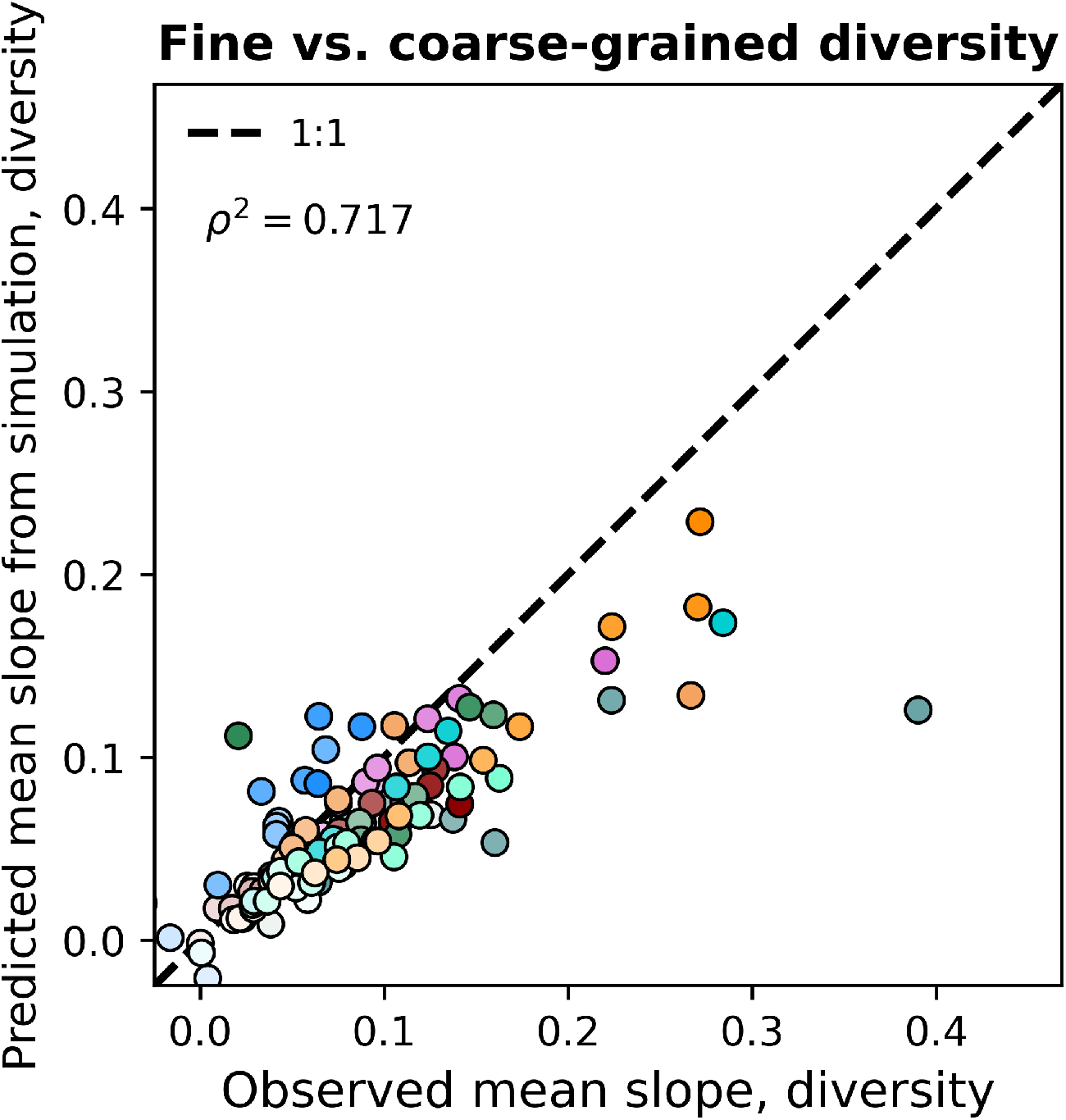
Including correlations allows the gamma distribution to capture observed diversity slopes. Observed fine vs. coarse-grained diversity slopes could be quantitatively reproduced under phylogenetic coarse-graining by simulating correlated gamma distributed AFDs at the OTU-level. The colorscale used here is identical to the colorscale used in Fig. 3. Squared Pearson correlation coefficients (*ρ*^2^) are computed over all slopes for all taxa across all coarse-graining scales.

## Discussion

The results of this study demonstrate that macroecological patterns in microbial communities remain largely invariant across taxonomic and phylogenetic scales. By focusing on the predictions of the SLM, an interaction-free model of microbial growth under environmental fluctuations, we were able to evaluate the extent that measures of biodiversity can be predicted under coarse-graining. We were largely able to predict said measures using the same model with parameters estimated from data across scales, implying that certain macroecological patterns of microbial communities remained self-similar across taxonomic and phylogenetic scales. Building off of this result, we investigated the dependence of community measures between different degrees of coarse-graining, a pattern that has been formalized as the Diversity Begets Diversity hypothesis [14, 15]. The prediction derived from the sampling form of the gamma distribution quantitatively captured the observed slopes of the fine vs. coarse-grained relationship for richness, while it failed to capture the slope of diversity. By introducing correlations between abundance fluctuation distribution we were able to recover the slope of the fine vs. coarse-grained diversity relationship.

Our richness results complement past work demonstrating that occupancy, the constituent of richness, is highly dependent on two parameters: sampling depth (i.e., total read count) and the mean abundance of a community member [40]. Our ability to predict the relationship between fine and coarse-grained measures of richness using the gamma distribution, despite our inability to predict the variance of richness, suggest that correlations driving the slope of the fine vs. coarse-grained relationship is primarily driven by the effects of finite sampling. This past work, and the relationships between the mean abundance and occupancy evaluated in this manuscript, demonstrate that occupancy alone is unlikely to contain ecological information that is not already captured by the distribution of abundances across sites (i.e., the AFD). Our analyses of the relationship between fine and coarse-grained richness support this conclusion, as predictions derived from a gamma distribution quantitatively captured the observed slope. The success of an interaction-free model in predicting the slope of the fine vs. coarse-grained relationship is an indictment of the appropriateness of estimators that rely solely on the presence of a community member for identifying novel macroecological patterns, a measure that has been used to bolster support for the DBD hypothesis at the level of 16S rRNA amplicons as well as strains [14, 63]. Rather, estimates of richness harbor little information about the dynamics of a community across taxonomic and phylogenetic scales that is not already captured by the sampling form of the gamma distribution. Contrasting with richness, the predictions of diversity from the gamma distribution were unable to capture fine vs. coarse-grained relationships from empirical data. Given that measures of diversity incorporate information about the richness and evenness of a community [64], the comparative deficiency of our predictions for fine vs. coarse-grained diversity suggests that forms of the SLM that neglect interactions between community members cannot capture coarse-graining relationships that depend on the evenness of the distribution of abundances.

Macroecological patterns are not imbued with mechanistic explanation [65]. Rather, the onus is on the investigator to identify plausible mechanisms. Often in ecology this task is made easier by evaluating whether a model lacking a particular mechanism is capable of producing the observed pattern, that is, identifying an appropriate null. The novelty of the fine vs. coarse-grained relationship was previously assessed using a null model which assumed demographic equivalence among community members and community dynamics driven by demographic noise (i.e., the UNTB) [14, 32]. Empirical patterns of microbial abundance cannot be reasonably captured by such models, making predictions obtained from the UNTB invalid for evaluating the novelty of microbial macroecological patterns. In contrast, models that combine self-limiting growth with environmental noise reproduce several empirical patterns, making the SLM an appropriate choice for evaluating the novelty of fine vs. coarse-grained relationships [40, 44]. This is not a trivial detail, as there is historical precedence on the need to identify an appropriate null in order to investigate how fine and coarse-grained measures of biodiversity relate to one another, as one of the earliest adoptions of null model analysis in ecology was done to investigate the ratio of species to genera in a community [66, 67].

In this study, the predictions of the sampling form of the gamma distribution considerably improved when correlations between community members were included. This result suggests that rather than exclusively pointing to niche construction as previously suggested [14], any ecological mechanism that can capture the observed distribution of correlation coefficients is a plausible candidate. Given that models of consumer-resource dynamics have succeeded in capturing macroecological patterns [68, 69], including quantitatively predicting the distribution of correlation coefficients [11], it is reasonable to suggest that such mechanisms are ultimately responsible for the relationship between fine and coarse-grained measures of diversity and can be reduced to phenomenological models such as the SLM. Indeed, experimental investigations of the slopes evaluated here have found the existence of positive slopes in artificial communities maintained in a laboratory setting, where the strength of the correlation between fine and coarse-grained scales is driven by the secretion of secondary metabolites [9]. This mechanism, known as cross-feeding, can be viewed as compatible with the concept of niche construction [16] as well as with the original interpretation of Madi et al. [14].

In the interest of providing macroecological insight into the DBD hypothesis, we solely focused on coarse-graining procedures that relied on phylogenetic reconstruction and taxonomic assignment. However, it is worth noting that it is also possible to coarse-grain community members by the strength of their correlations (i.e., sum the abundances of each pair of community members with the strongest correlation in AFDs). This procedure has been named the phenomenological renormalization group method due to its ability to identify if and where a system is stable despite knowing little about the system’s dynamics (i.e., fixed points in nonlinear systems) [70, 71]. However, given that the AFD correlation between two community members is often inversely related to their phylogenetic distance, such an analysis would likely be redundant, as coarse-graining based on the strength of correlation would effectively coarse-grain the most closely related community members [54].

A major goal of this study was to evaluate the novelty of macroecological patterns that were used to bolster support for the DBD hypothesis. We used the same dataset in order to ensure generality and commensurability with past research efforts. However, it is worth inspecting how the use of a global survey dataset constrains the inferences one can make. Throughout this study we implicitly assumed that an ensemble approach is valid, meaning that we viewed different sites/hosts as virtual copies of a given environment. This assumption is likely valid for time-series studies where the distribution of microbial abundances remains stationary with respect to time [72], as the stationary solution of the SLM has successfully characterized microbial community time-series at both the level of OTUs [40] and strains [49]. Given these past results, we predict that the fine vs. coarse-grained relationship results presented here will remain valid in longitudinal studies where community members fluctuate around a stationary with respect to time.

## Materials and methods

### Data acquisition and processing

To ensure that our analyses were generalizable across ecosystems and commensurate with prior DBD investigations, we used amplicon sequence data from the V4 region of the 16S rRNA gene generated and curated by the Earth Microbiome Project [1, 14]. We restricted our analysis to the quality control (QC)-filtered subset of the EMP, which was annotated using the closed-reference database SILVA [73] and consists of 96 studies culminating in 23,828 total samples with each processed sample having ≥10,000 reads. We downloaded the public Silva reference tree for OTUs with 97% similarity `97_otus.tre` from the EMP database. We identified nine heavily sampled environments in the metadata file `emp_qiime_mapping_qc_filtered.tsv` and selected 100 random sites from each environment. Summary statistics for each environment can be found in the Supporting Information (Tables S1, S2).

We briefly note that our occupancy and richness predictions depend on the form of the gamma distribution that explicitly accounts for sampling as a multinomial process. The multinomial distribution describes the probability of sampling *n* reads given a relative abundance of *x* and total read count *N* with replacement, a process we can model as the Poisson limit of a binomial sampling process for individual community members. Given this choice and the past success of the gamma distribution, we deviated from past analyses by electing to not sub-sample read counts to the same depth, as the process of sampling without replacement would bias the sampling distribution for rare community members [14].

### Coarse-graining protocol

Taxonomic coarse-graining was performed as the summation of the abundances of all OTUs within a given taxonomic group. We removed taxa with indeterminate labels to prevent potential biases due to taxonomic misassignment, (e.g., “uncultured”, “ambiguous taxa”, “candidatus”, “unclassified”, etc.). Manual inspection of EMP taxonomic annotations revealed a low number of OTUs that had been assigned the taxonomic label of their host (e.g., *Arachis hypogaea* (peanut)). These marked OTUs were removed from all downstream analyses.

Phylogenetic coarse-graining was performed using the phylogenetic tree provided by SILVA 123 `97_otus.tre` in the EMP release. Each internal node of a phylogenetic tree was collapsed if the mean branch lengths of its descendants was less than a given distance. All phylogenetic operations were performed using the Python package `ETE3` [74].

### Deriving biodiversity measure predictions

While the gamma distribution as the stationary solution of the SLM and the sampling form of the gamma distribution have been previously derived [40], we briefly outline relevant derivations here for the convenience of the reader before deriving the predicted richness and diversity of a community. We define the SLM as the following Langevin equation

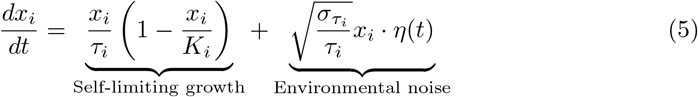

Here *τ*_*i*_, *K*_*i*_, and 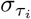 represent the timescale of growth, the carrying capacity, and the coefficient of variation of growth rate fluctuations, respectively. Multiplicative environmental noise is captured by the product of a linear frequency term, the coefficient of variation of growth rate fluctuations, and a Brownian noise term *η*(*t*) that introduces stochasticity into the equation. The expected value of *η*(*t*) is *η*⟨(*t*)⟩ = 0 [75].

The dependence of *η*(*t*′) at time *t*′ on an earlier time *η*(*t*) is defined as ⟨*η*(*t*)*η*(*t*′)⟩ = *δ*(*t*−*t*′) [75]. This standard definition means that if the noise term is shifted in time, it has zero correlation with itself. We briefly note that because DBD patterns were originally investigated in Madi et al. using an ensemble of sites that belong to the same type of ecosystem rather than the timeseries of a single site [14], the gamma distribution alone does not prove the validity of the SLM nor does it prove alternatively formulated stochastic differential equations of ecology that also predict a gamma distribution (e.g., [76]). However, given that the SLM has successfully characterized the temporal dynamics of microbial communities, we believe that this model is an appropriate formulation for investigating DBD patterns [40, 44, 49].

In contrast to the SLM, macroecological predictions can be derived from the UNTB. There are many forms of the UNTB, but the novelty of observed fine vs. coarse-grained relationships was assessed using a form of the UNTB that predicts that the distribution of community member abundances within a given site follows a zero-sum multinomial distribution [14, 32]. For the convenience of the reader the predicted richness using the form of the UNTB relevant to this study has been rederived (Supporting Information).

The stationary distribution of the SLM can be derived using the Itô ↔ Fokker-Planck equivalence and solving for the stationary solution [40, 77], resulting in the gamma distributed AFD. Through the SLM, we can define the mean relative abundance and its squared inverse coefficient of variation as 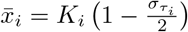 and 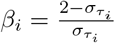, respectively. These are parameters that were estimated from the empirical data and were used below to obtain predictions. Using these definitions and the stationary distribution of Eq. 5, we obtained the gamma distribution

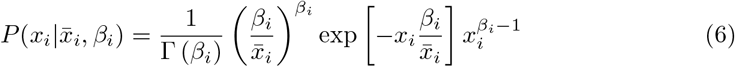

When we sequence microbial communities, one obtains read counts rather than actual abundances. Therefore, it is necessary to account for the reality of sampling when we apply Eq. 6 to empirical data. We can account for sampling by first assuming that the probability of observing a single community member can be modeled as a binomial sampling process. Given that the total number of reads is typically large (*N* ≫ 1) and the typical relative abundance of a community member is much smaller than one (*x*_*i*_≪1), the binomial can be approximated as a Poisson sampling process with the following probability of sampling *n* reads

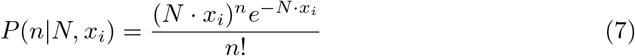

This formulation of the sampling process is convenient, as it can be used to obtain an analytic solution for the probability of observing *n* reads given 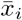 and *β*_*i*_, the parameters we estimate from the data. This distribution can be obtained solving the convolution of the Poisson and the gamma distribution [40]. The resulting distribution can be considered a negative binomial distribution if sites have identical sampling depths [78]. Using this distribution, we calculated the probability of obtaining *n*_*m*_ reads out of a total sampling depth of *N*_*m*_ for the *i*th OTU in sample *m* as

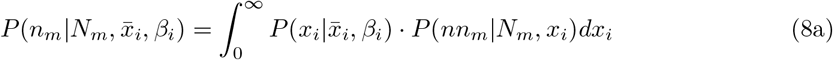

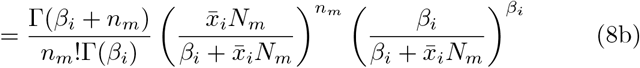

This distribution requires two parameters that can be estimated from the data (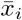 and *β*_*i*_) and one parameter that is known (total number of reads, *N*_*m*_). This equation will be used to obtain predictions of measures of biodiversity. First, noticing that the probability of a community member’s absence is the complement of its presence, we can define the expected occupancy of a community member across *M* sites as

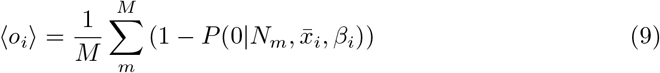

And the second moment of occupancy as

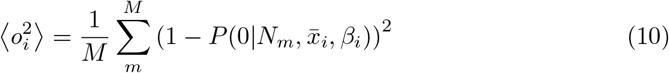

from which we defined the predicted variance of occupancy

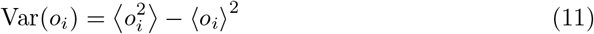

The success of our predictions was assessed using the relative error.

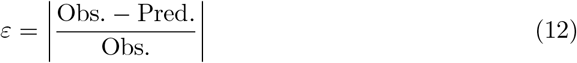

Using the definition of occupancy from the sampling form of the gamma distribution, we derived the expected richness of a community as

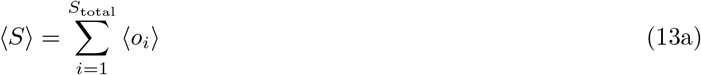

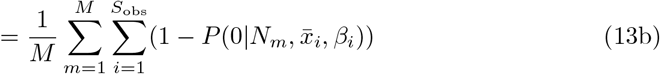

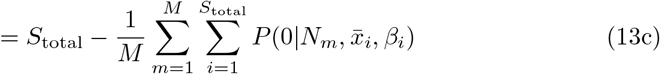

where *S*_total_ is the total number of observed community members. Similarly, we derived the expected value of Shannon’s diversity [64].

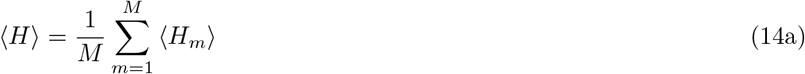

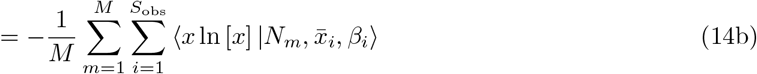

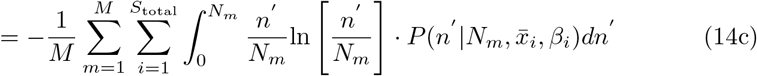

In physics parlance, these predictions neglect interactions between community members, also known as mean-field predictions. We then calculated the mean-field prediction of Eq. 13 from empirical data. However, there is no known analytic solution for the integral inside the sum of Eq. 14. To calculate ⟨*H*⟩, we performed numerical integration on each integral for each taxon in each sample at a given coarse-grained resolution using the `quad()` function from `SciPy`.

To predict the variance of each measure we derived the expected value of the second moment, assuming independence among community members. We derived the second moments of richness and diversity

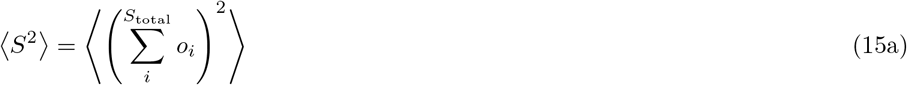

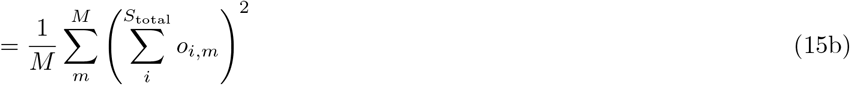

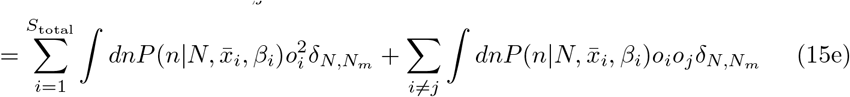

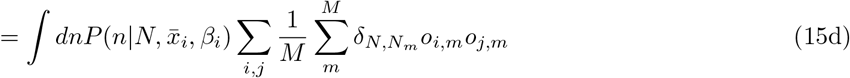

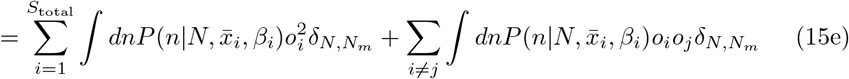

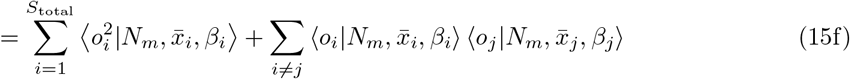

where *δ*_*i,j*_ is the Kronecker delta.

By performing an analogous series of operations, we obtained the expected value of the second moment for diversity.

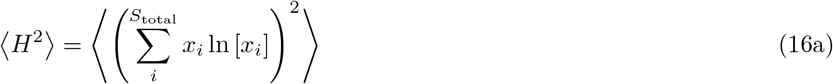

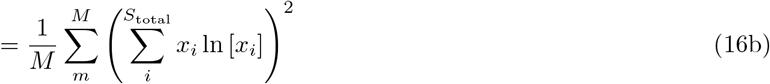

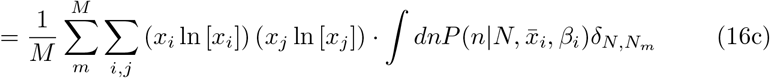

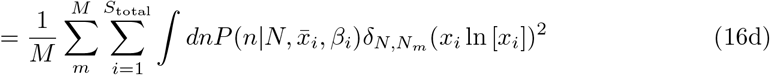

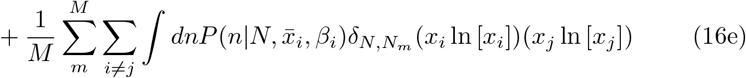

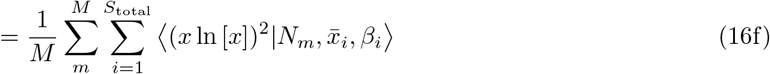

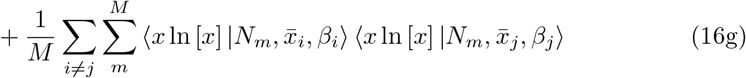

Where the expected value of the second moment of the diversity term is defined as 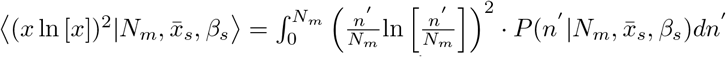.From which we obtained the expected value of the variance

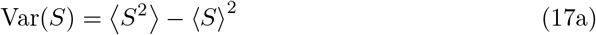

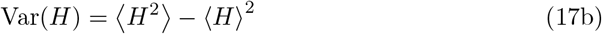

We predicted the mean and variance of richness and diversity separately at each coarse-grained scale. Specifically, we coarse-grained the empirical data, estimate 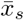 and *β*_*s*_ for each coarse-grained community member, and use these estimates to obtain a prediction for each measure of biodiversity.

It is worth noting why the above functions constitute predictions. To obtain values that we can compare with empirical data we estimated the mean and variance of relative abundance across sites for each community member at a given scale. These parameters were used to obtain the expected value of a community-level measure (e.g., richness) using a function. These functions were derived under the assumption that a given probability distribution (i.e., the gamma) provided an appropriate description of the distribution of relative abundances across sites. We then compared the expected value of a community-level measure to the mean value from empirical data and assessed the similarity between the two values.

### Fine vs. coarse-grained relationship slope inference

In order to predict the relationship between the measures within a coarse-grained group and that among all remaining groups, we calculated a vector of predicted richness or diversity estimates for all sites using Eq. 3a or Eq. 4a within a given coarse-grained group and 3b or Eq. 4b among the remaining groups. This “leave-one-out” procedure was originally implemented in Madi et al., where the authors examined the slope of fine vs. coarse-grained measures of diversity as a sliding window across taxonomic ranks with both the fine and coarse scales increasing with each rank (e.g., genus:family, family: order, etc.) [14]. To maintain consistency, we used the same definition for our predictions. We also extended the definition to the case of phylogenetic coarse-graining, where we compared fine and coarse scales using different phylogenetic distances while retaining the same ratio (e.g., 0.1:0.3, 0.3:0.5, etc.). Slopes were estimated using ordinary least squares regression with `SciPy`. Throughout the manuscript the success of a prediction was evaluated by calculating its relative error as follows: we only inferred the slope if a fine-grained group had at least five members. We only examined the slopes of a given coarse-grained threshold if at least three slopes could be inferred.

### Simulating communities of correlated gamma-distributed AFDs

Correlated gamma-distributed AFDs were simulated by performing inverse transform sampling. For each environment with *M* sites, an *M* × *S*_obs_ matrix **Z** was generated from the standard Gaussian distribution using the empirical *S*_obs_×*S*_obs_ correlation matrix calculated from relative abundances. The cumulative distribution **U** = Φ(**Z**)_Gaus._ was calculated and a matrix of the abundances of community members across sites was obtained using the point percentile function of the gamma distribution and the empirical distribution of mean relative abundances and the squared inverse coefficient of variation of abundances: 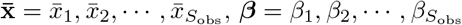. To simulate the process of sampling, each community of the resulting *M* × *S*_obs_ matrix of true relative abundances 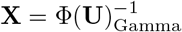 was sampled using a multinomial distribution with the empirical distribution of total read counts.

## Data and code availability

All sequencing data used in this study was obtained from the EMP (URL: ftp://ftp.microbio.me/emp/release1/). Processed data are available on Zenodo, DOI: `10.5281/zenodo.7692046`. All code written for this study is available on GitHub under a GNU General Public License: `macroeco phylo`

## Acknowledgments

This work was supported by the NSF Postdoctoral Research Fellowships in Biology Program under Grant No. 2010885 (W.R.S.).

## Author contributions

W.R.S. and J.G. conceptualized the project, completed the derivations, and wrote the manuscript. W.R.S. performed all analyses.

