## Supplemental Information for "Investigating macroecological patterns in coarse-grained microbial communities using the stochastic logistic model of growth"

#### UNTB richness predictions

Below we rederive a prediction for richness using the form of the UNTB used by Madi et al. as a point of comparison to the SLM predictions derived in the main manuscript (1; 2). When the size of a metacommunity tends towards an asymptotic limit, the stationary distribution for community members of relative abundance $x$ approaches the following continuous distribution (3)

$$P(x|\theta)dx = \frac{\theta}{x}(1-x)^{\theta-1}dx \tag{S1}$$

where $\theta$ is Hubbell's biodiversity parameter (also known as Fisher's $\alpha$ (4)).

Using this distribution, we can obtain an expression for the expected number of community members with $n$ sampled individuals out of a total sample size of $N$.

$$S(n|N,m,\theta) = \theta \int_0^1 P(n|N,m,x)\frac{(1-x)^{\theta-1}}{x}dx \tag{S2}$$

where $P(n;N,m,x)$ is the probability of sampling $n$ individuals of relative abundance $x$ given a total sample size $N$

$$P(n|N,m,x) = \binom{N}{n}\frac{\Gamma(n+\gamma x)}{\Gamma(\gamma x)}\frac{\Gamma(N+\gamma(1-x)-n)}{\Gamma(\gamma(1-x))}\frac{\Gamma(\gamma)}{\Gamma(\gamma+N)} \tag{S3}$$

where $\gamma = \frac{m(N-1)}{1-m}$. The function Eq. S2 is known as the migration-limited zero-sum multinomial distribution (ZSM) (2). As $m \to 1$, Eq. S2 approaches a limiting form known as the metacommunity zero-sum multinomial distribution (mZSM). The process of sampling community members under the mZSM can be represented as a binomial distribution.

$$S(n|N,\theta) = \theta \int_0^1 x^N(1-x)^{N-n}\frac{(1-x)^{\theta-1}}{x}dx \tag{S4}$$

Similar to our analysis using the SLM, the binomial can be approximated as a Poisson distribution

$$S(n|N,\theta) = \theta \int_0^1 e^{-xN}\frac{(xN)^n}{n!}\frac{(1-x)^{\theta-1}}{x}dx \tag{S5}$$

The resulting integral can be obtained by using the change of variable $y = xN$ and rearranging terms, then approximating the upper limit of integration as infinity (since $N \gg 1$)

$$S(n|N,\theta) = \theta \int_0^1 e^{-y} \frac{(y)^n}{n!} \left(1 - \frac{y}{N}\right)^{\theta-1} \frac{N}{y} \frac{dy}{N} \tag{S6a}$$

$$= \frac{\theta}{n} \int_0^1 \underbrace{e^{-y} \frac{y^{n-1}}{(n-1)!}}_{\text{Gamma distribution}} \left(1 - \frac{y}{N}\right)^{\theta-1} dy \tag{S6b}$$

$$\approx \frac{\theta}{n} \int_0^\infty \underbrace{e^{-y} \frac{y^{n-1}}{(n-1)!}}_{\text{Gamma distribution}} \left(1 - \frac{y}{N}\right)^{\theta-1} dy \tag{S6c}$$

$$= \frac{\theta}{n} \left\langle \left(1 - \frac{Y}{N}\right)^{\theta-1} \right\rangle \tag{S6d}$$

By rearranging terms we obtain a gamma distribution with shape parameter $n$ and rate parameter 1. Because we integrated over a product with a gamma distribution, the variable $Y$ is a gamma distributed random variable. We can then expand term in the integral using a Taylor series around $Y = n$ and by noticing that $\langle Y \rangle = n$ and $\langle Y^2 \rangle = n^2 + n$ under a gamma distribution

$$S(n|N,\theta) = \frac{\theta}{n} \left(1 - \frac{n}{N}\right)^{\theta-1} + \frac{1}{2} \frac{\theta(\theta-1)(\theta-2)}{N^2} \left(1 - \frac{n}{N}\right)^{\theta-3} + \mathcal{O}\left(N^{-3}\right) \tag{S7}$$

We can then predict the richness of a community by summing the abundances from 1 to $N$

$$S(N,\theta) = \sum_{n=1}^{N} S(n|N,\theta) \tag{S8}$$

This quantity represents the total observed richness of a sample from a panmictic infinite metacommunity after accounting for sampling. Predictions of mean richness over $M$ sites can then be calculated as

$$\langle S(\theta) \rangle = \frac{1}{M} \sum_{m=1}^{M} S(N_m, \theta) \tag{S9}$$

#### Simulating fine vs. coarse-grained richness slopes under the UNTB

We followed the procedure in Madi et al. to obtain fine vs. coarse-grained slopes for richness so that they could be compared to predictions obtained from the SLM (5). We simulated SADs according the mZSM model outlined above using the `rmzsm()` function from the R package `sads` v0.4.2. We simulated 100 SADs using the empirical distribution of total read counts and the total number of observed OTUs. We set the biodiversity parameter $\theta = 50$ for all environments. The SADs returned by `rmzsm()` contain no zeros, meaning that values of richness are identical for all UNTB SADs. In order to introduce zeros so that richness estimates could vary, we followed the procedure used in Madi et al. where each simulated SAD was rarefied to 5,000 individuals. We repeated this rarefaction procedure on the empirical SADs. We then performed taxonomic and phylogenetic coarse-graining and fine vs. coarse-grained slope inference using the procedure described in the Materials and methods.

### 1 Figures

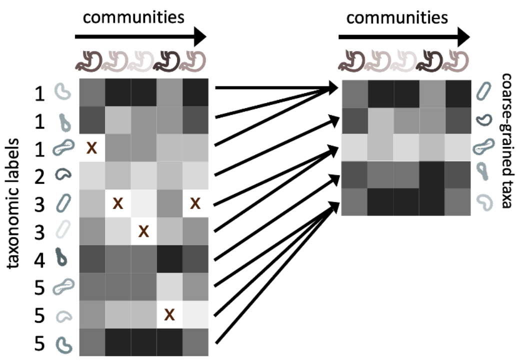


**Figure S1. The process of coarse-graining using taxonomic information.** Taxonomic assignment in 16S rRNA amplicon sequence data provides the opportunity to investigate how properties of communities vary at different taxonomic scales. A straightforward means of coarse-graining is to sum the abundances of OTUs/ASVs that belong to the same taxonomic group.

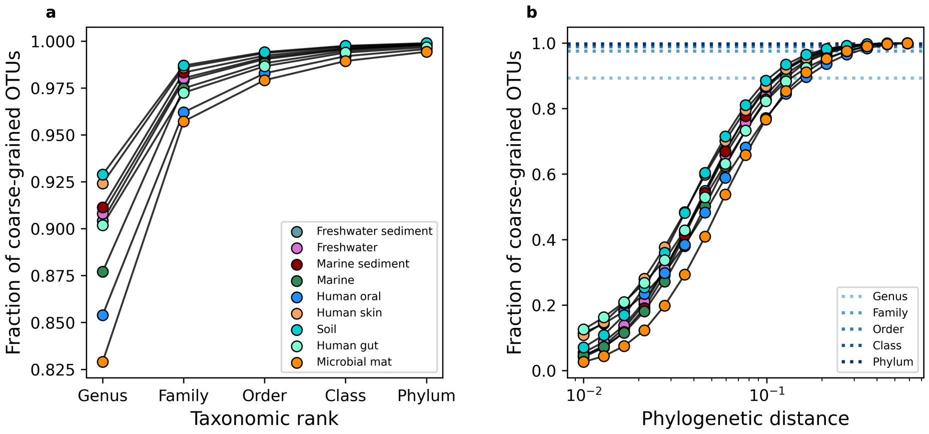


**Figure S2. Examining the change in relative richness under coarse-graining.** To gain an intuition for how the number of community members changes in the face of coarse-graining, we can examine the fraction of OTUs that remain across scales of **a)** taxonomic and **b)** phylogenetic coarse-graining. In **b** the mean fraction over environments for a given taxonomic rank is plotted for reference.

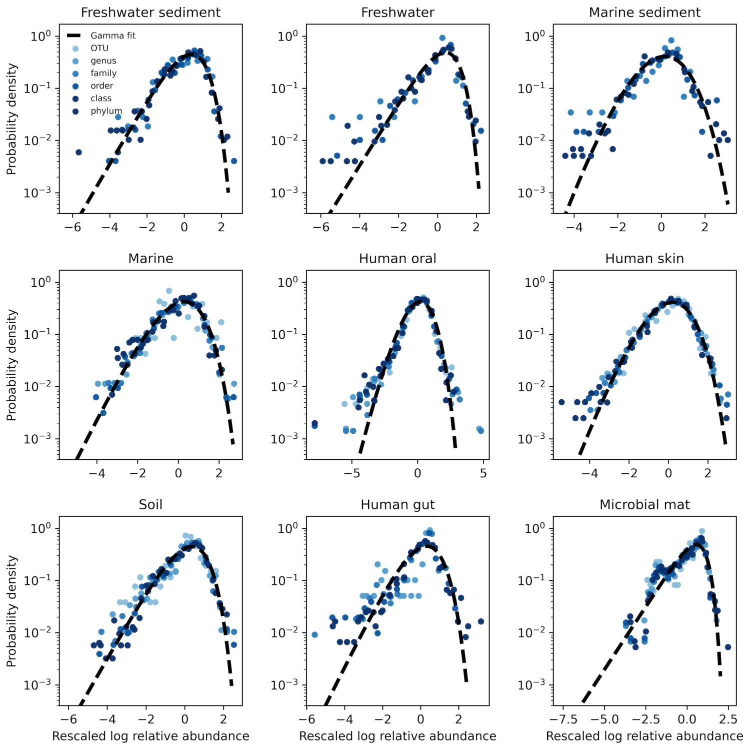


**Figure S3. The AFD of all environments under taxonomic coarse-graining.** To control for the effect of sampling we only examined the AFDs of OTUs that were present in all sites (i.e., an occupancy of one). The single exception was the microbial mat, where we used a minimum occupancy of 0.75 as the environment harbored no OTUs that were present among all sites. Black lines represent fits of the gamma distribution obtained using `SciPy`.

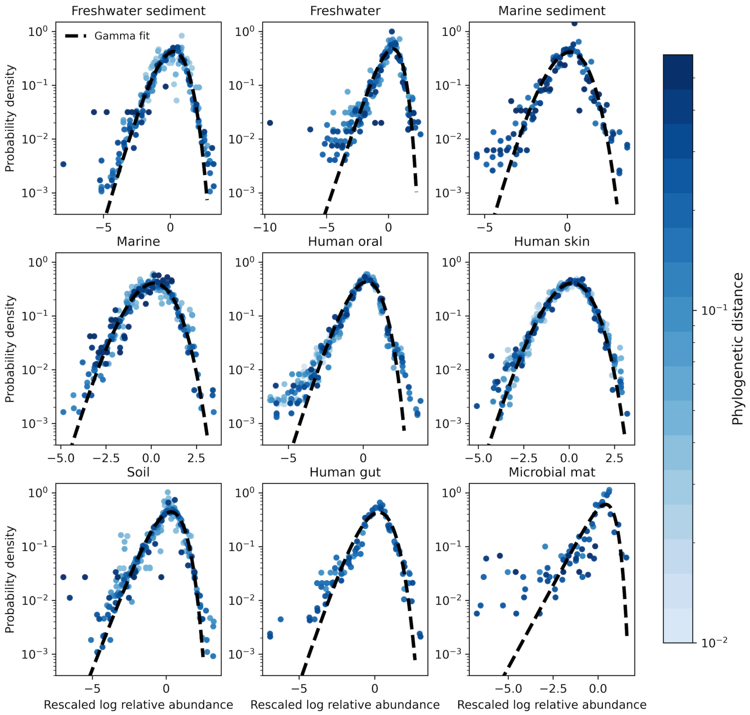


**Figure S4. The AFD of all environments under phylogenetic coarse-graining.** To control for the effect of sampling we only examined the AFDs of OTUs that were present in all sites (i.e., an occupancy of one). Black lines represent fits of the gamma distribution obtained using `SciPy`.

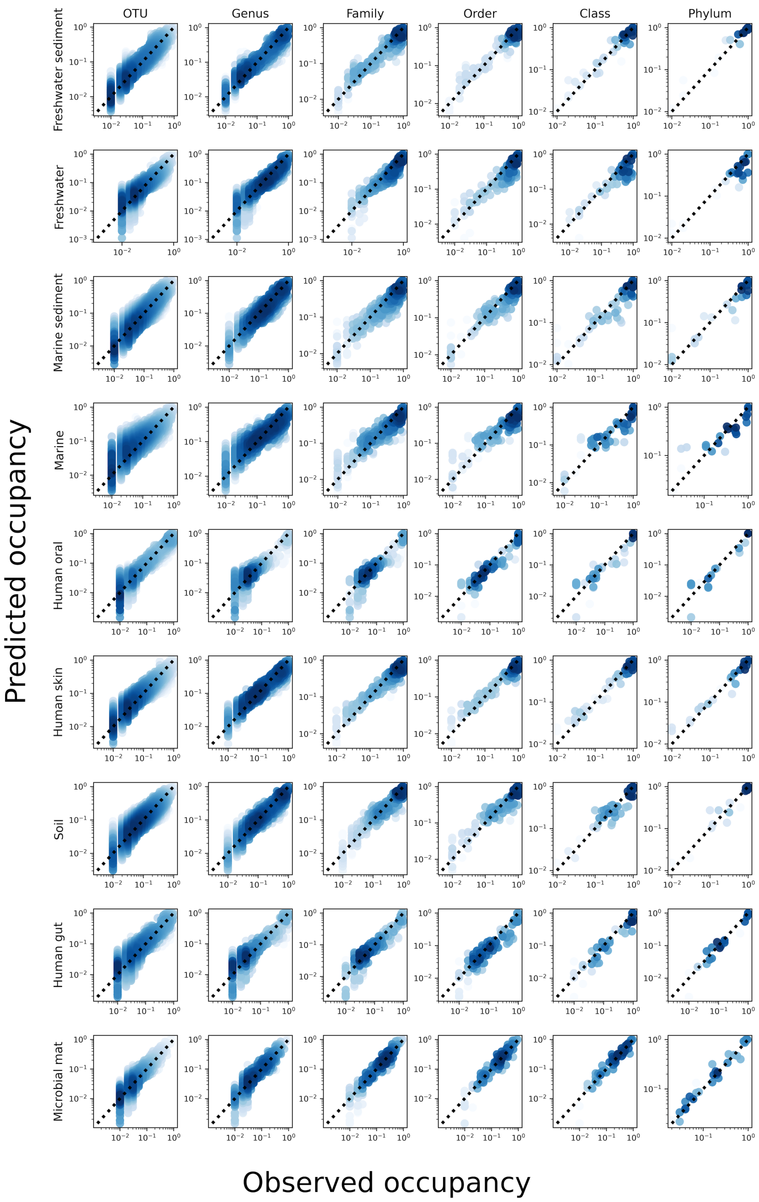


**Figure S5.** The predicted occupancy across sites for a gamma distributed AFD under taxonomic coarse-graining for all environments.

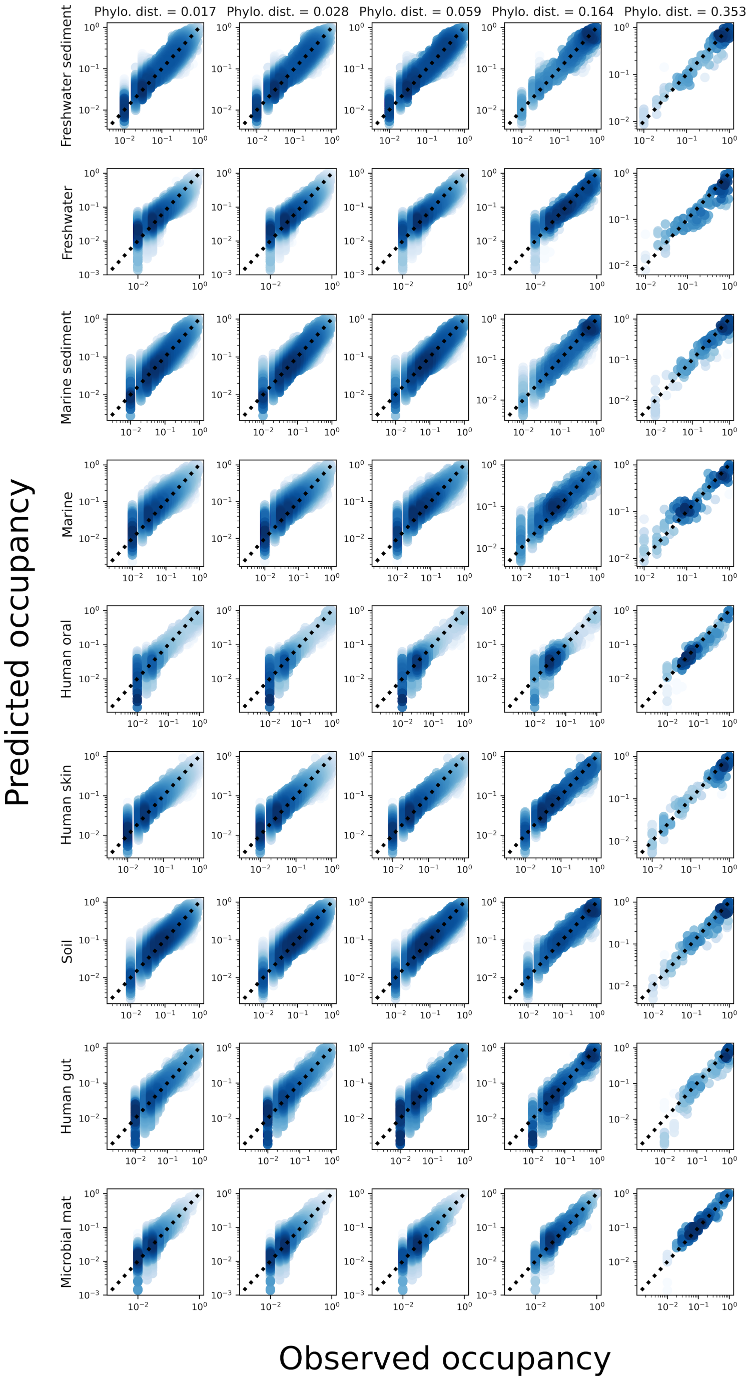


**Figure S6.** The predicted occupancy across sites for a gamma distributed AFD under phylogenetic coarse-graining for all environments.

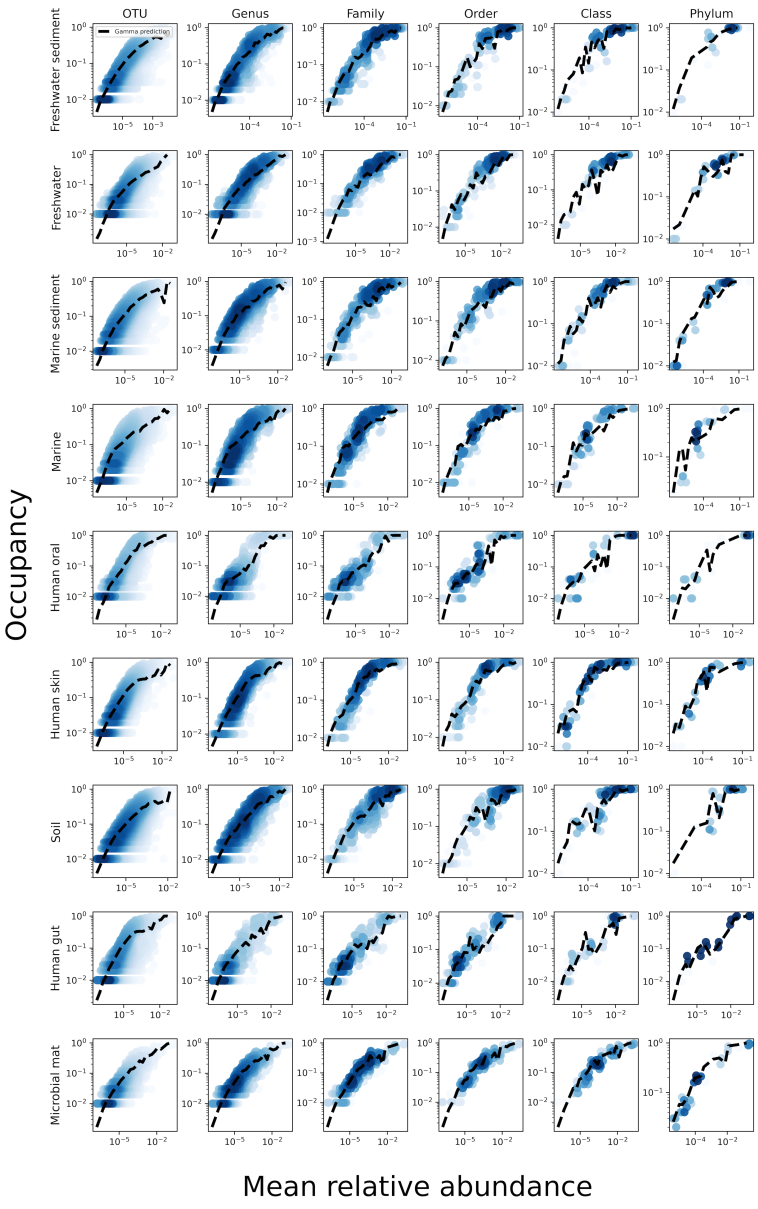


**Figure S7.** The relationship between the mean abundance across sites and the occupancy for various taxonomic coarse-graining scales. The black line represents the prediction of the gamma distribution.

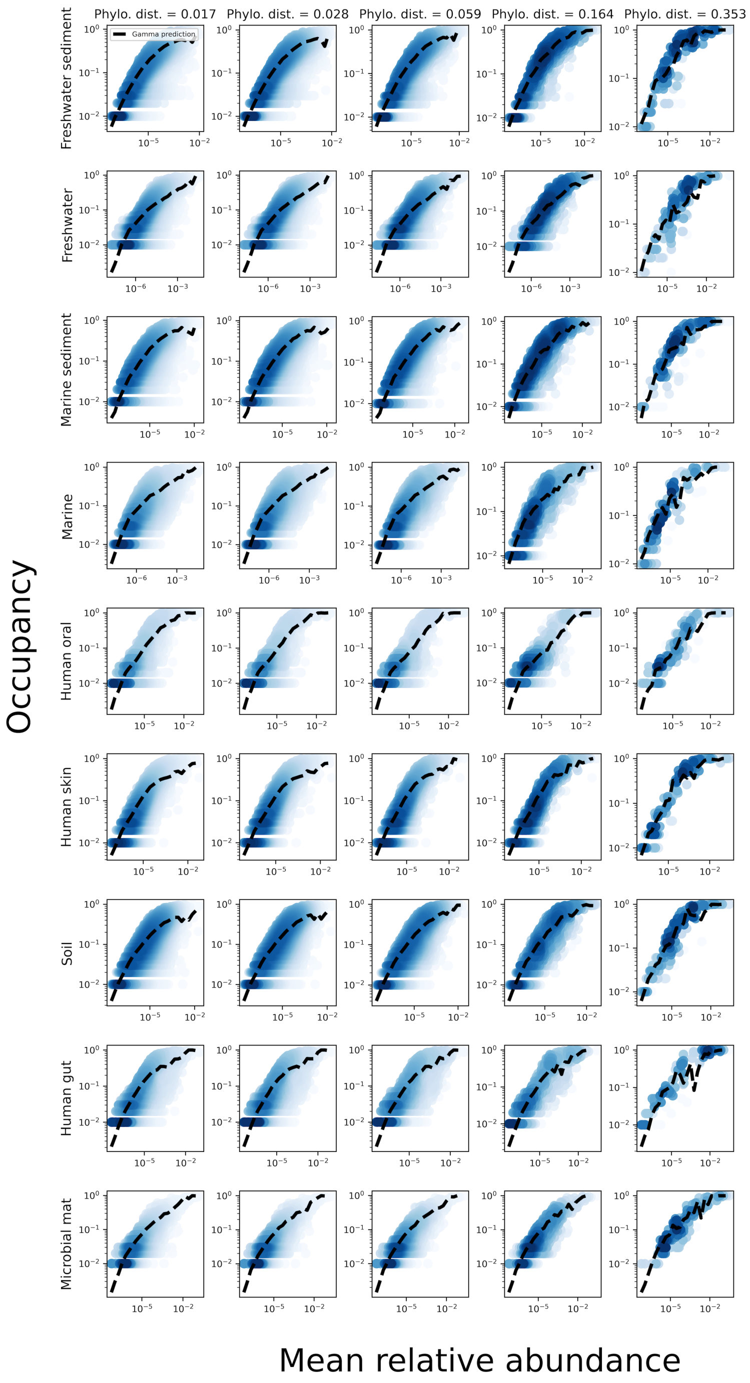


**Figure S8.** The relationship between the mean abundance across sites and the occupancy for various phylogenetic coarse-graining scales. The black line represents the prediction of the gamma distribution.

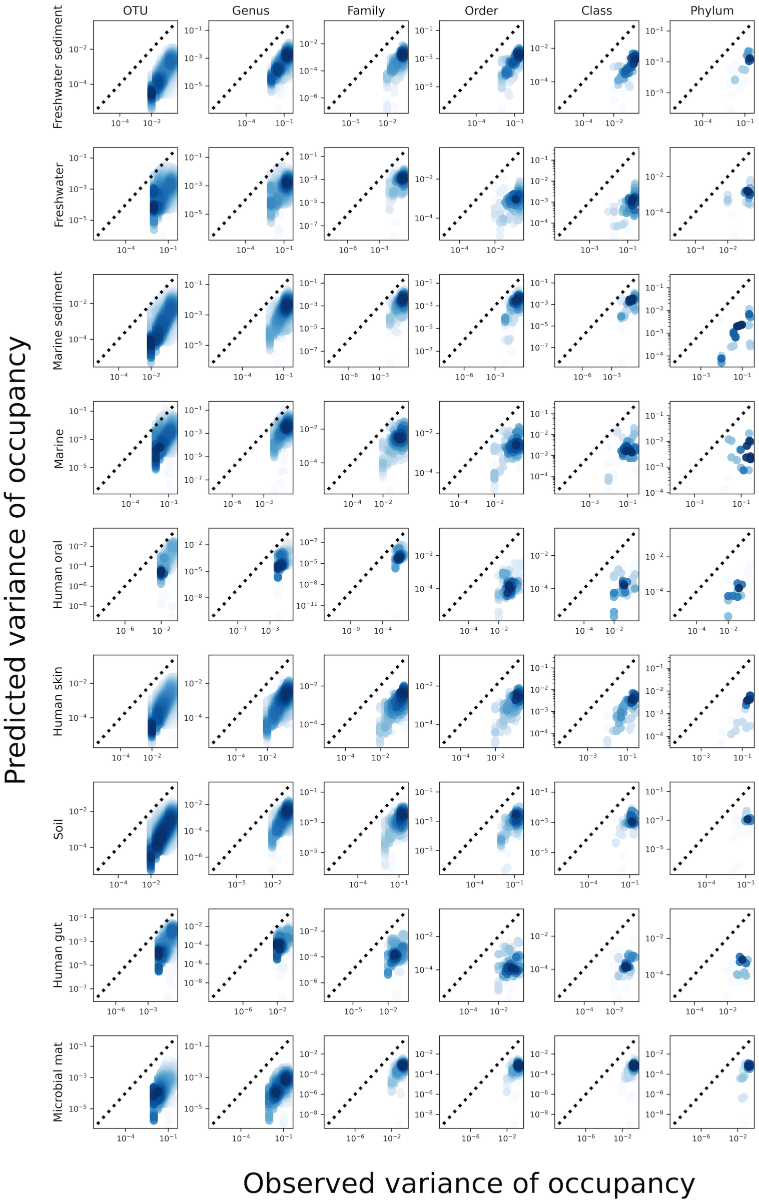


**Figure S9. Predictions of the variance of occupancy failed across taxonomic coarse-graining thresholds.** While the gamma distribution succeeded in predicting mean occupancy, it failed to predict the variance of occupancy.

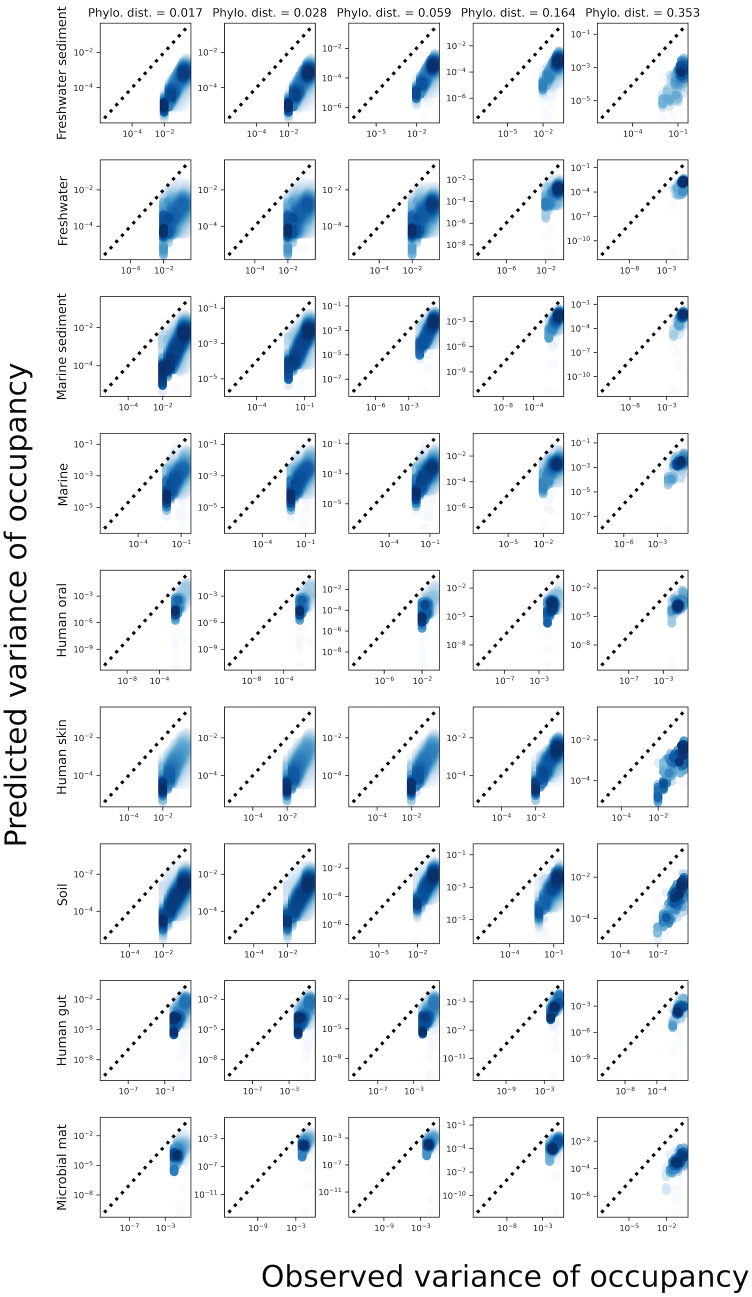


**Figure S10. Predictions of the variance of occupancy failed across phylogenetic coarse-graining thresholds.** Analogous plot of S9 for phylogenetic coarse-graining.

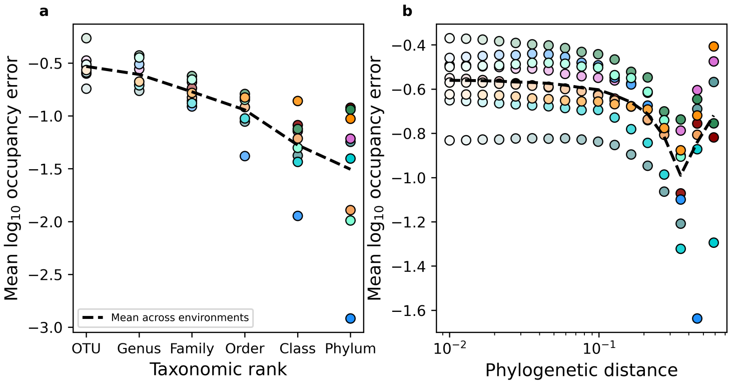


**Figure S11. Occupancy predictions of the gamma remained invariant despite coarse-graining.** Despite **a)** taxonomic and **b)** phylogenetic coarse-graining the mean relative error of occupancy predictions using the gamma did not increase. Instead, the error tended to decline over extended coarse-graining scales, only increasing for phylogenetic coarse-graining when communities were coarse-grained to $\lesssim 5$ members.

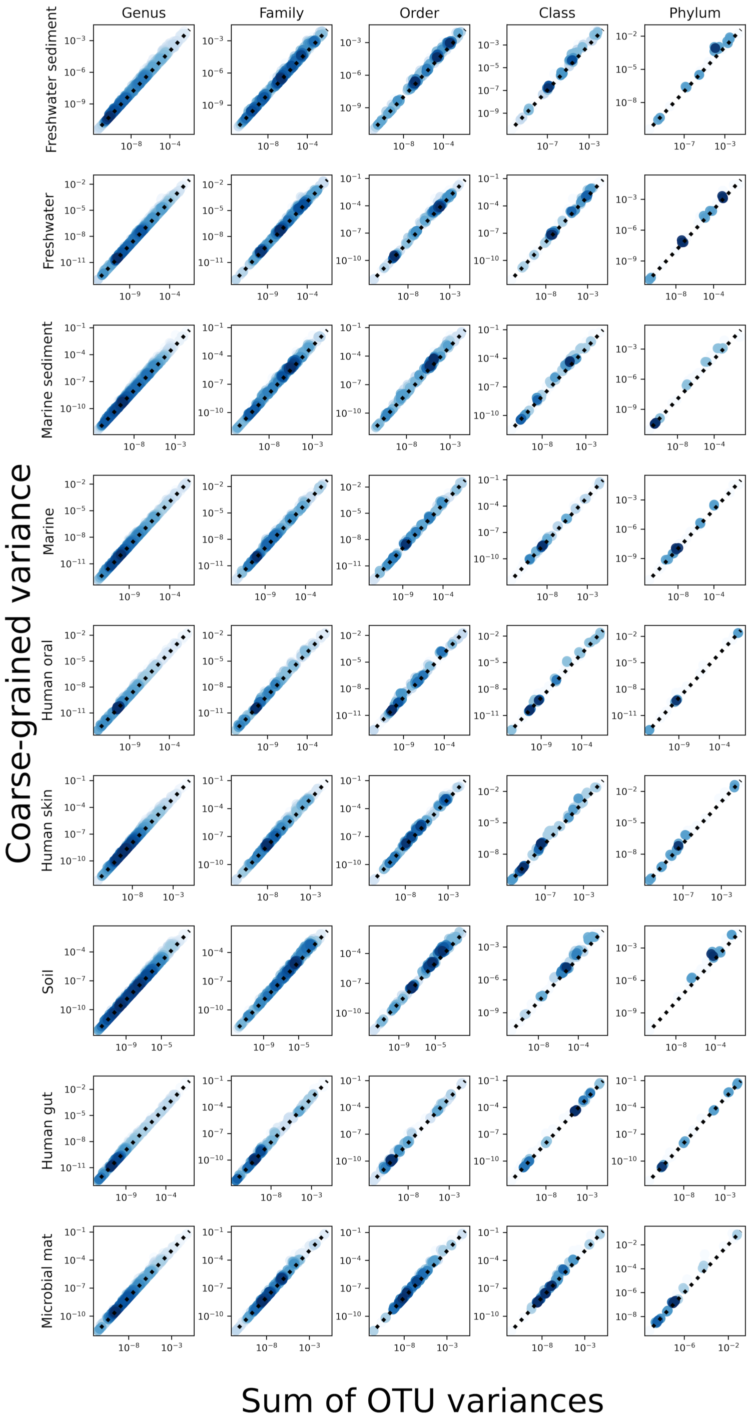


**Figure S12.** The sum of the variances of OTUs was close to the value of the variance of a taxonomic coarse-grained group, implying that the contribution of covariance to the variance of a given coarse-grained group was low.

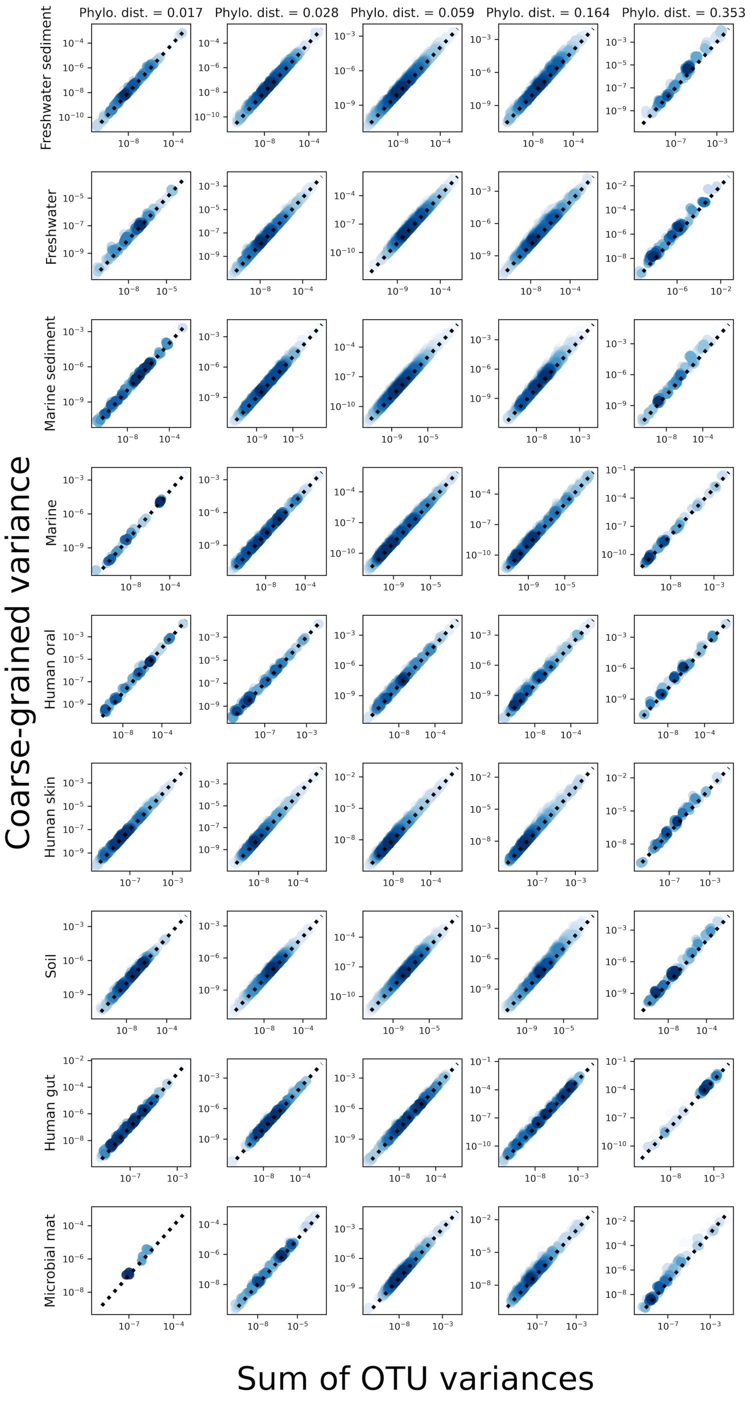


**Figure S13.** The analysis presented in Fig. S12 but for phylogenetic coarse-graining.

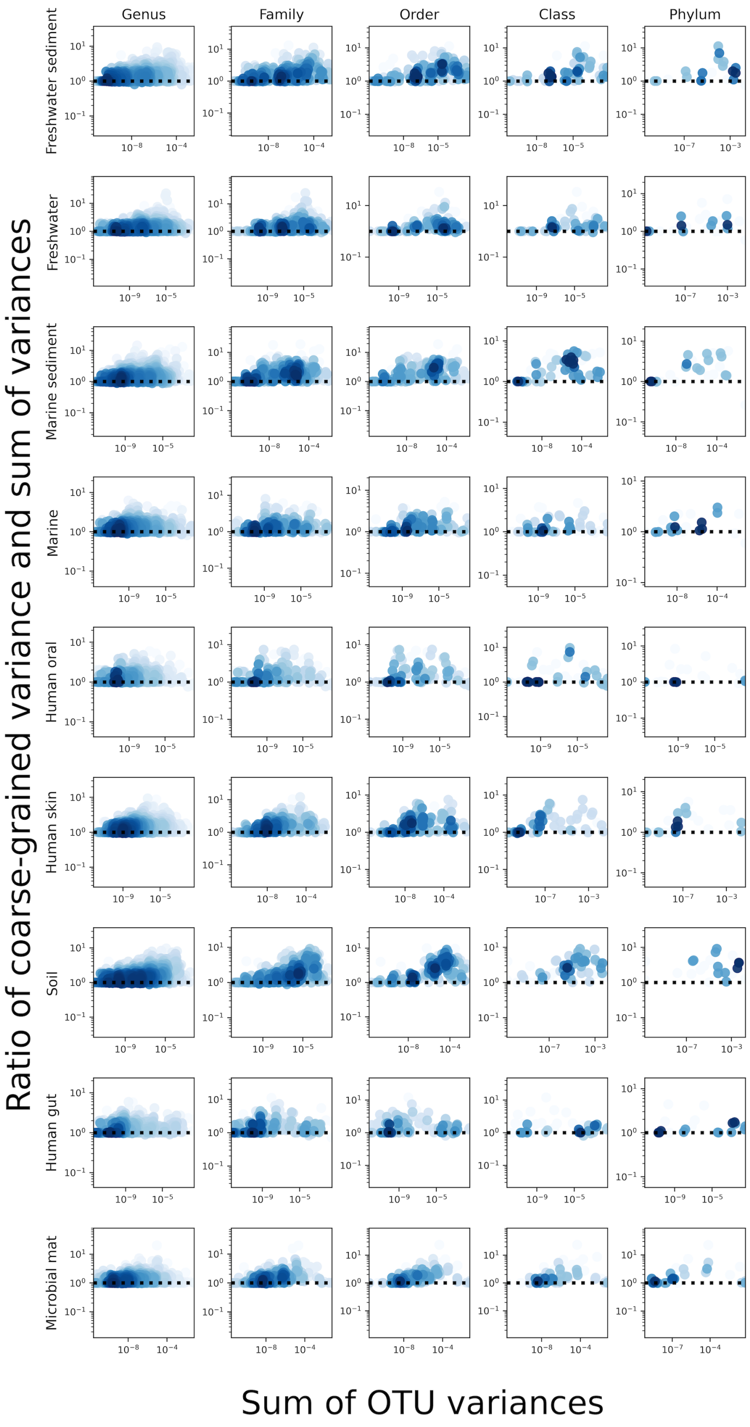


**Figure S14.** The plot presented in Fig. S12 but with the ratio of coarse and fine-grained variances plotted on the y-axis for the purpose of visualising deviations from the 1:1 line.

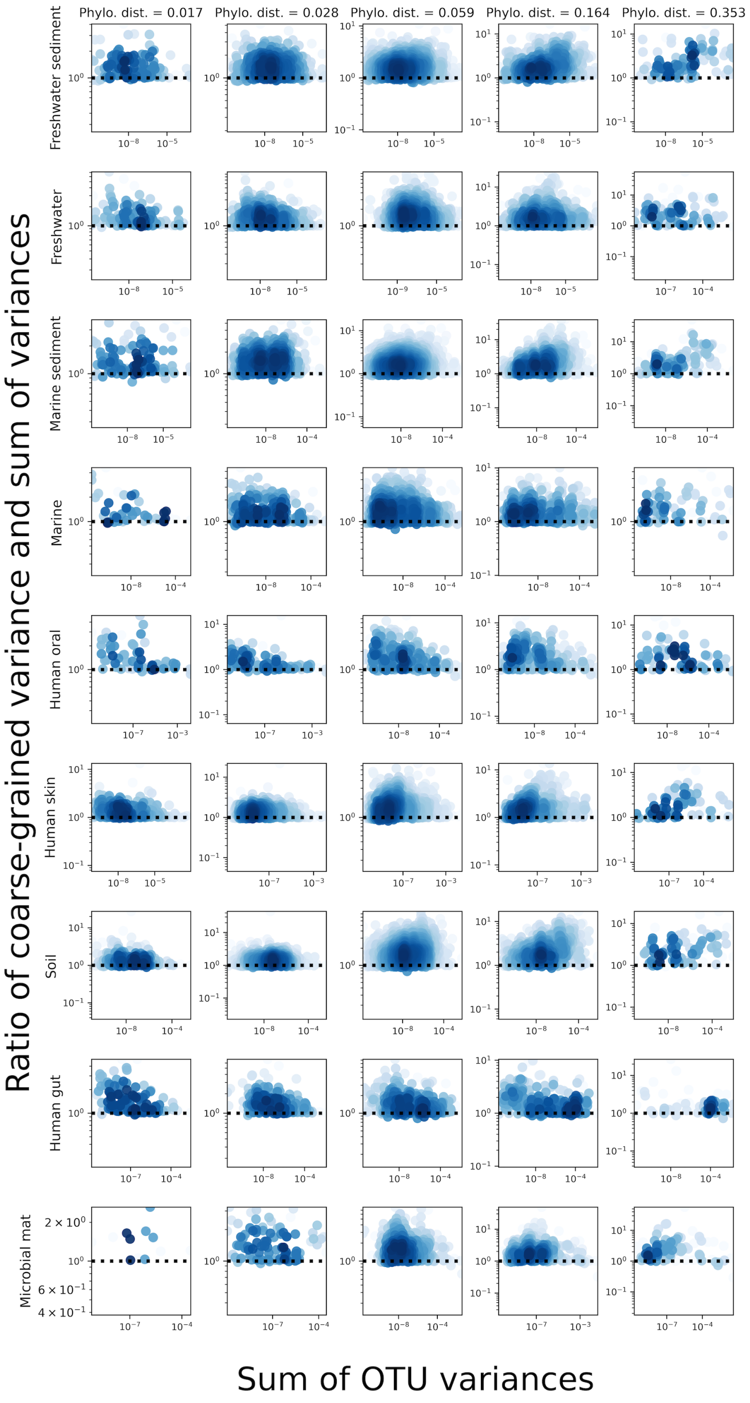


**Figure S15.** The analysis presented in Fig. S14 but for phylogenetic coarse-graining.

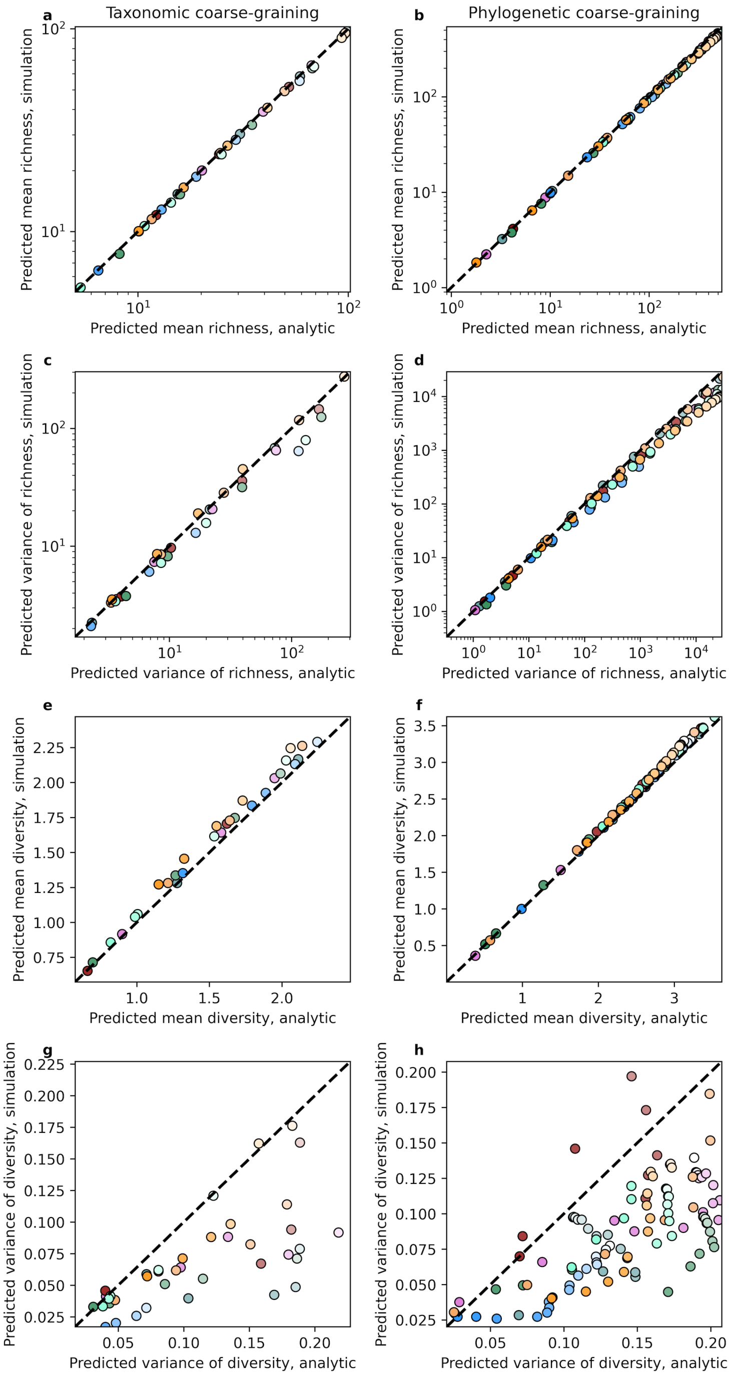


**Figure S16.** The analytic predictions of the mean and variance of richness and diversity versus the results of simulations that assume gamma distributed AFDs and reads drawn from a multinomial distribution.

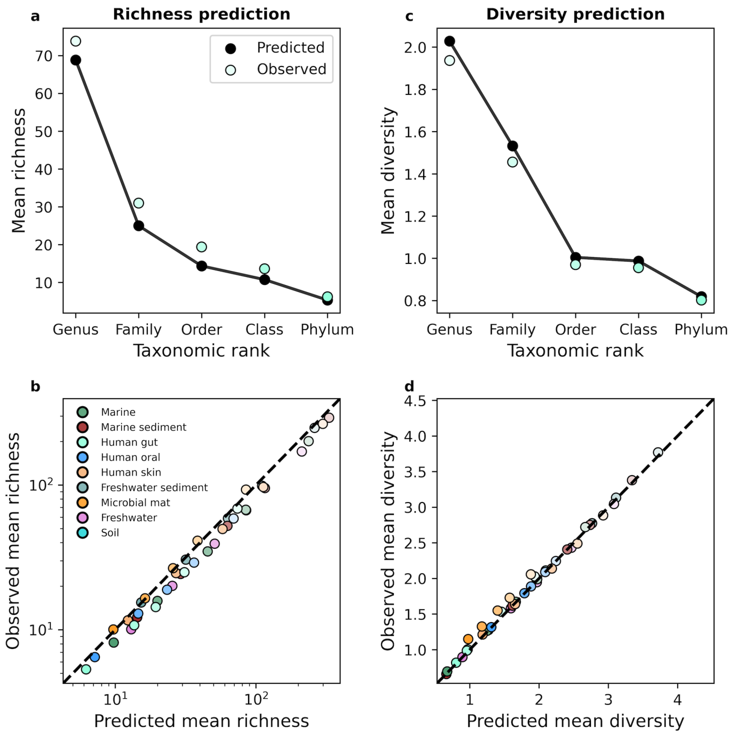


**Figure S17. The gamma distribution successfully predicted mean richness and diversity under taxonomic coarse-graining.** An equivalent set of analyses as depicted in Fig. 3 for taxonomic coarse-graining.

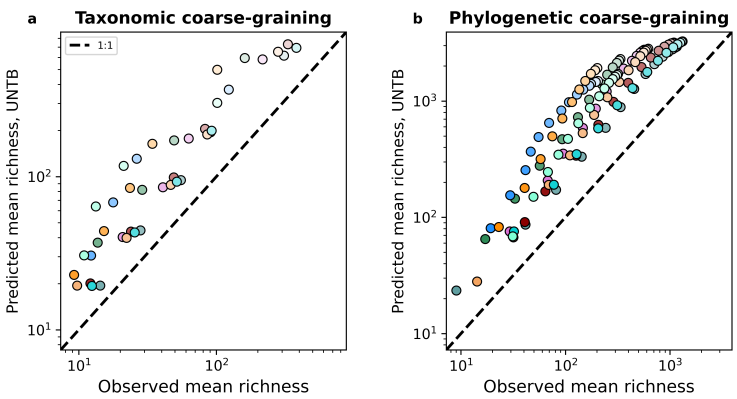


**Figure S18. UNTB failed to predict mean richness.** UNTB consistently overpredicted richness under both taxonomic and phylogenetic coarse-graining.

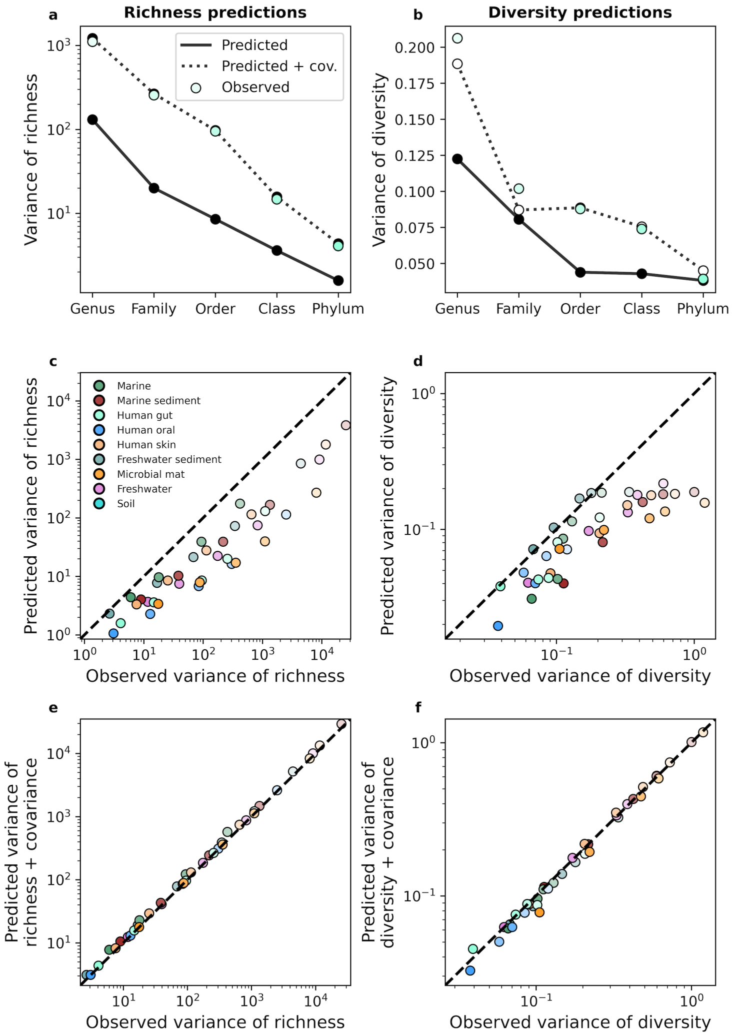


**Figure S19. A gamma AFD can only predict the variance of richness and diversity under taxonomic coarse-graining when covariance is included.** An equivalent set of analyses as depicted in Fig. 4 for taxonomic coarse-graining.

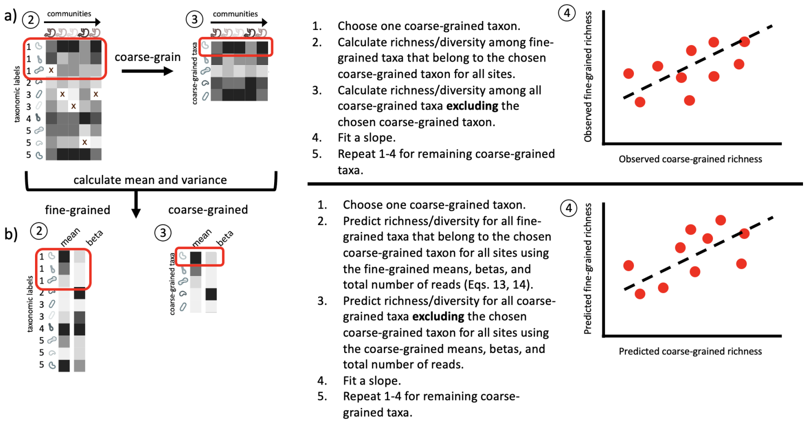


**Figure S20.** Conceptual diagram illustrating how fine vs. coarse-grained slopes are inferred. **a)** Slopes were inferred from empirical data by estimating fine and coarse-grained measures of biodiversity (for this diagram, richness) according to the leave-one-out procedure used by Madi et al. (5). **b)** The mean relative abundance and beta (squared inverse coefficient of variation) were then estimated from the data and then used to predict fine and coarse-grain estimates of a given measure of biodiversity using the same leave-one-out procedure. Separate regressions were then fit to the empirical and predicted fine and coarse-grained measures of biodiversity.

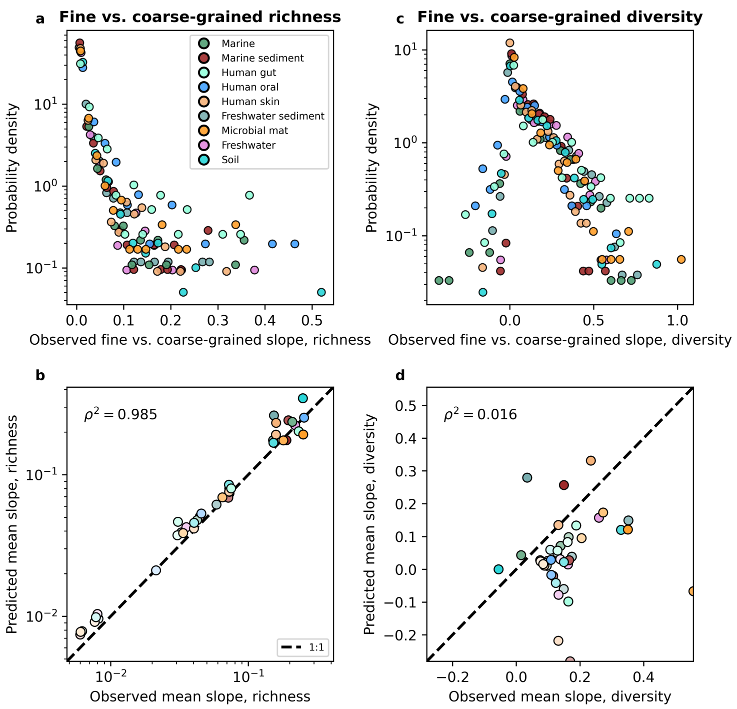


**Figure S21. The gamma distribution as a tool for investigating the novelty of fine vs. coarse-grained slopes.** An equivalent set of analyses as depicted in Fig. 5 for taxonomic coarse-graining.

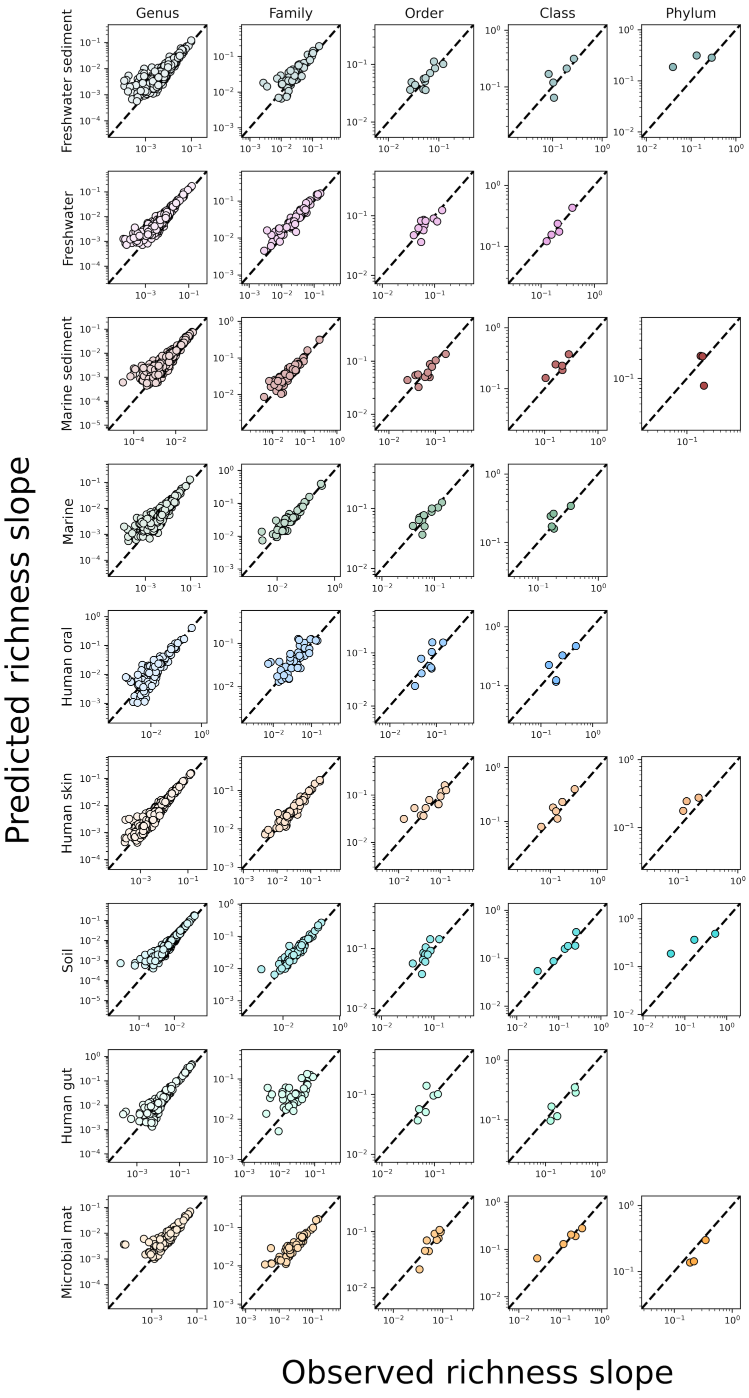


**Figure S22.** The predicted slopes of fine vs. coarse-grained richness from the sampling form of the gamma distribution under taxonomic coarse-graining.

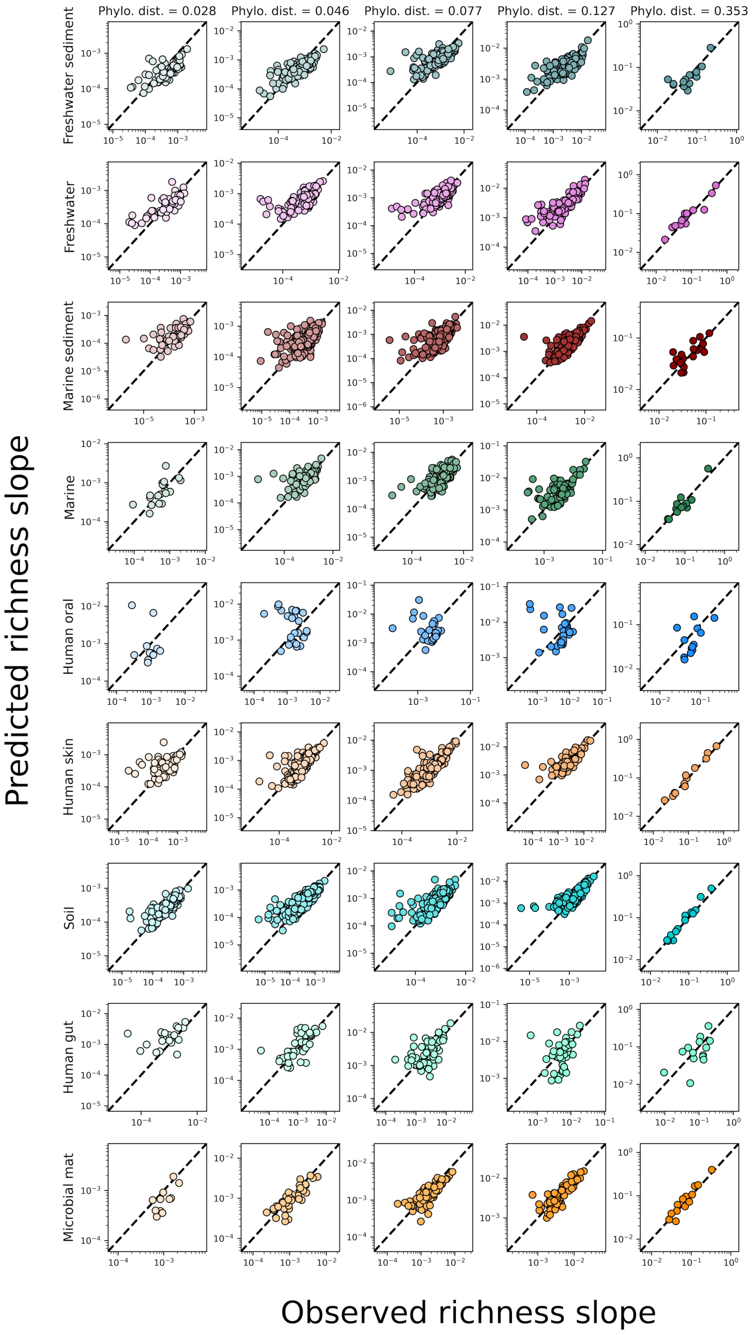


**Figure S23.** The predicted slopes of fine vs. coarse-grained richness from the sampling form of the gamma distribution under phylogenetic coarse-graining.

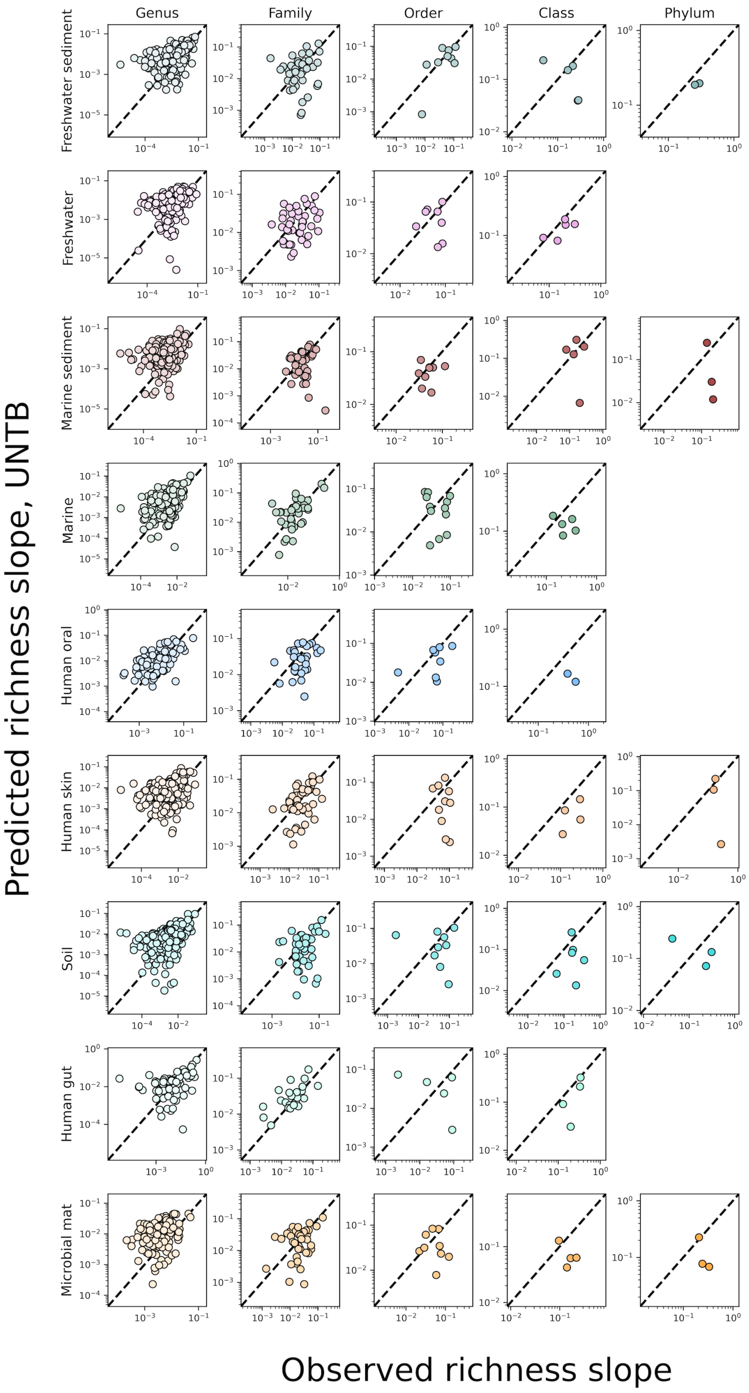


**Figure S24.** The predicted slopes of fine vs. coarse-grained richness under taxonomic coarse-graining using the UNTB.

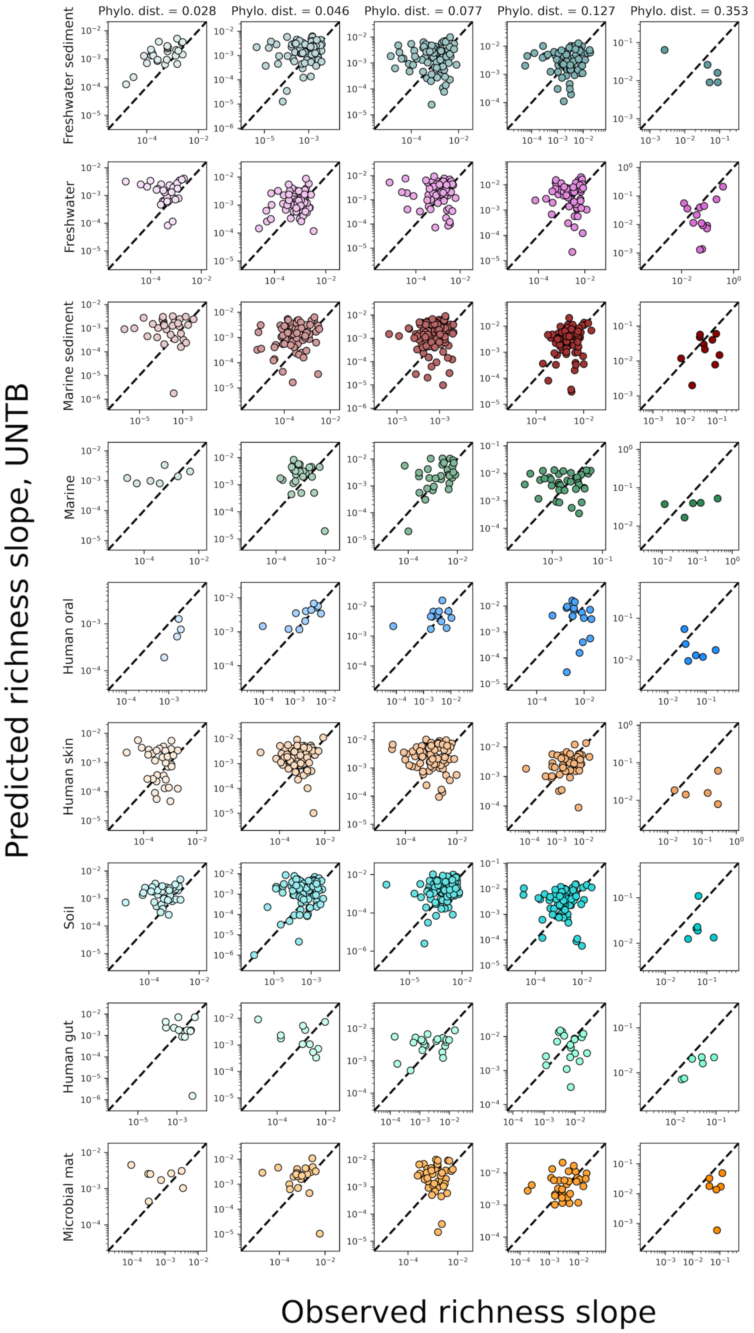


**Figure S25.** The predicted slopes of fine vs. coarse-grained richness under phylogenetic coarse-graining using the UNTB.

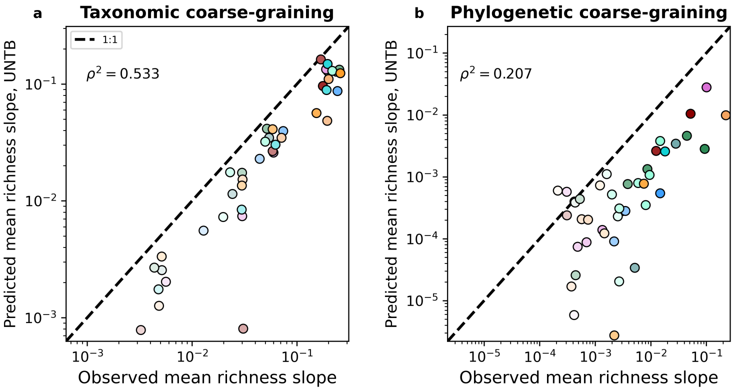


**Figure S26.** The mean predicted slopes of fine vs. coarse-grained richness under **a)** taxonomic and **b)** phylogenetic coarse-graining using the UNTB.

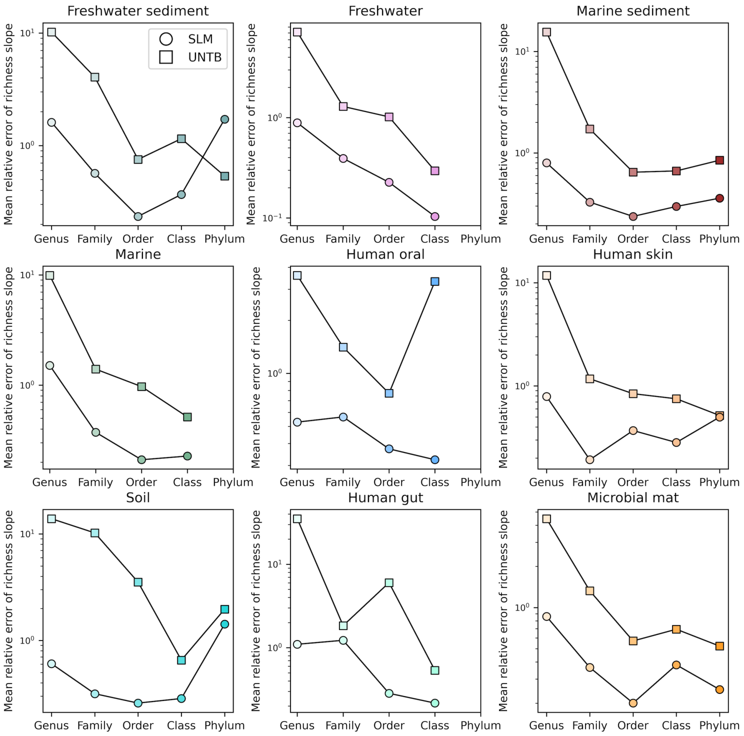


**Figure S27.** Comparisons of the relative error of fine vs. coarse-grained richness slope predictions between the SLM and UNTB for taxonomic coarse-graining.

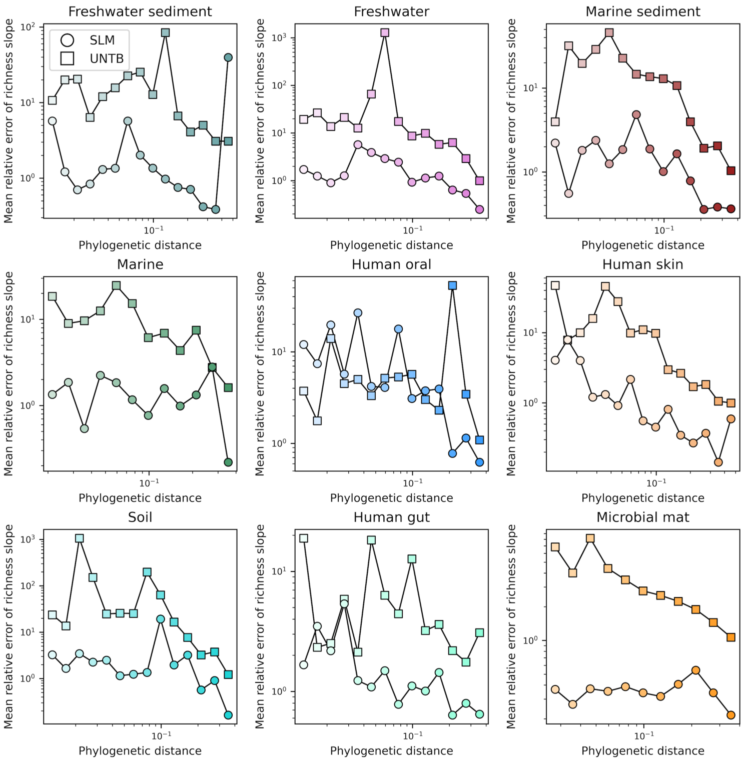


**Figure S28.** Comparisons of the relative error of fine vs. coarse-grained richness slope predictions between the SLM and UNTB for phylogenetic coarse-graining.

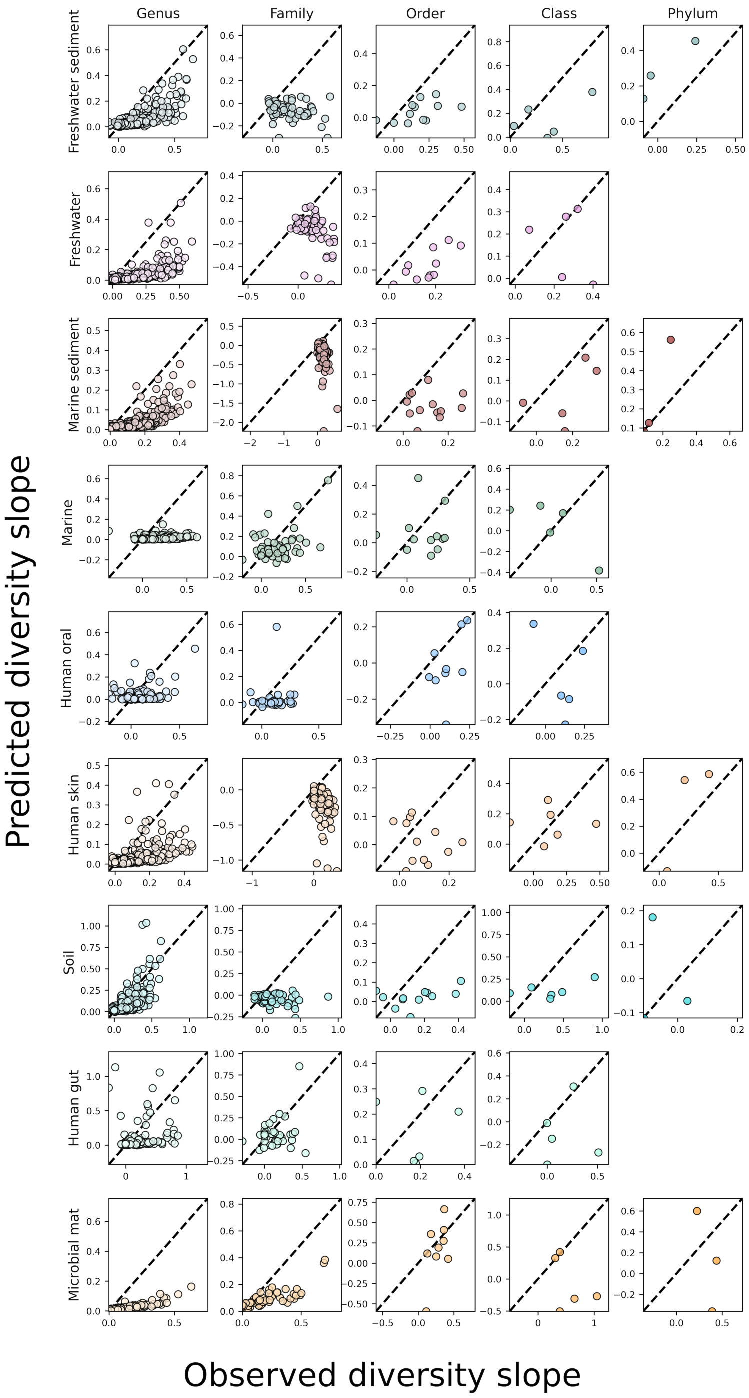


**Figure S29.** The predicted slopes of fine vs. coarse-grained diversity from the sampling form of the gamma distribution under taxonomic coarse-graining.

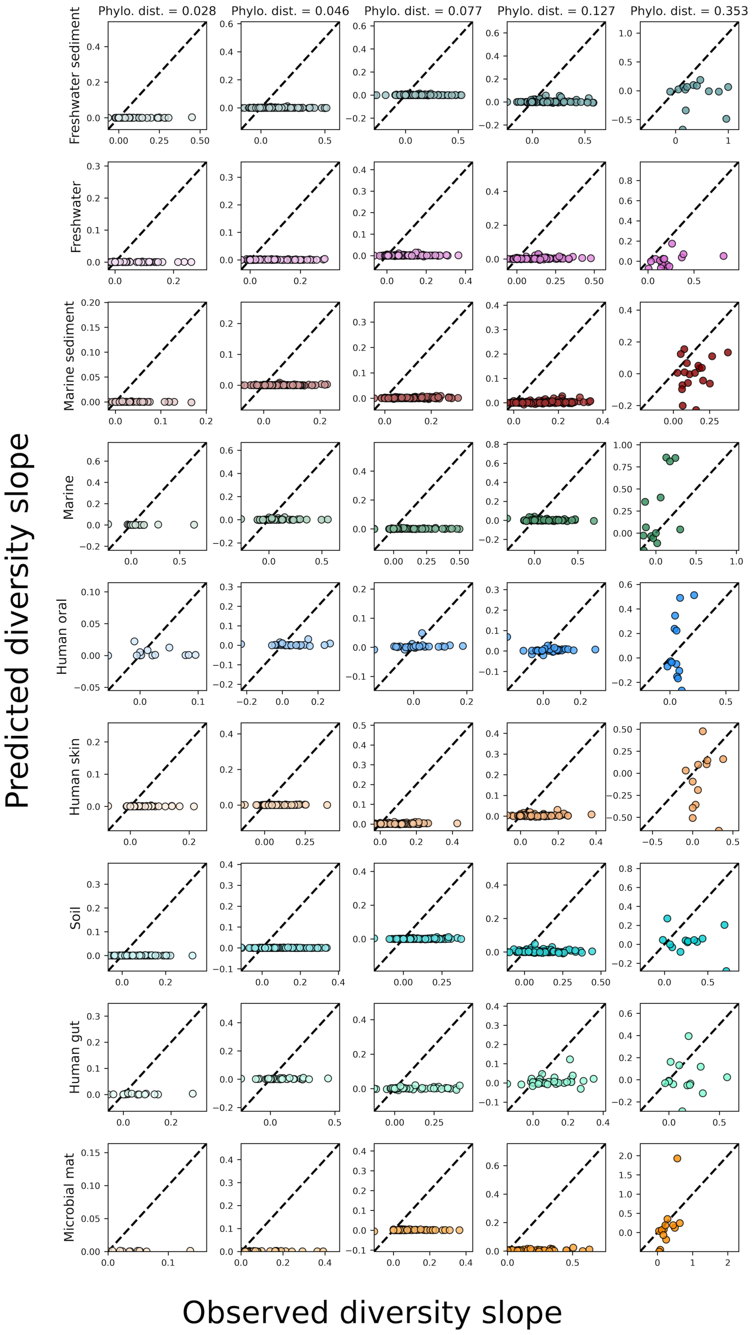


**Figure S30.** The predicted slopes of fine vs. coarse-grained diversity from the sampling form of the gamma distribution under phylogenetic coarse-graining.

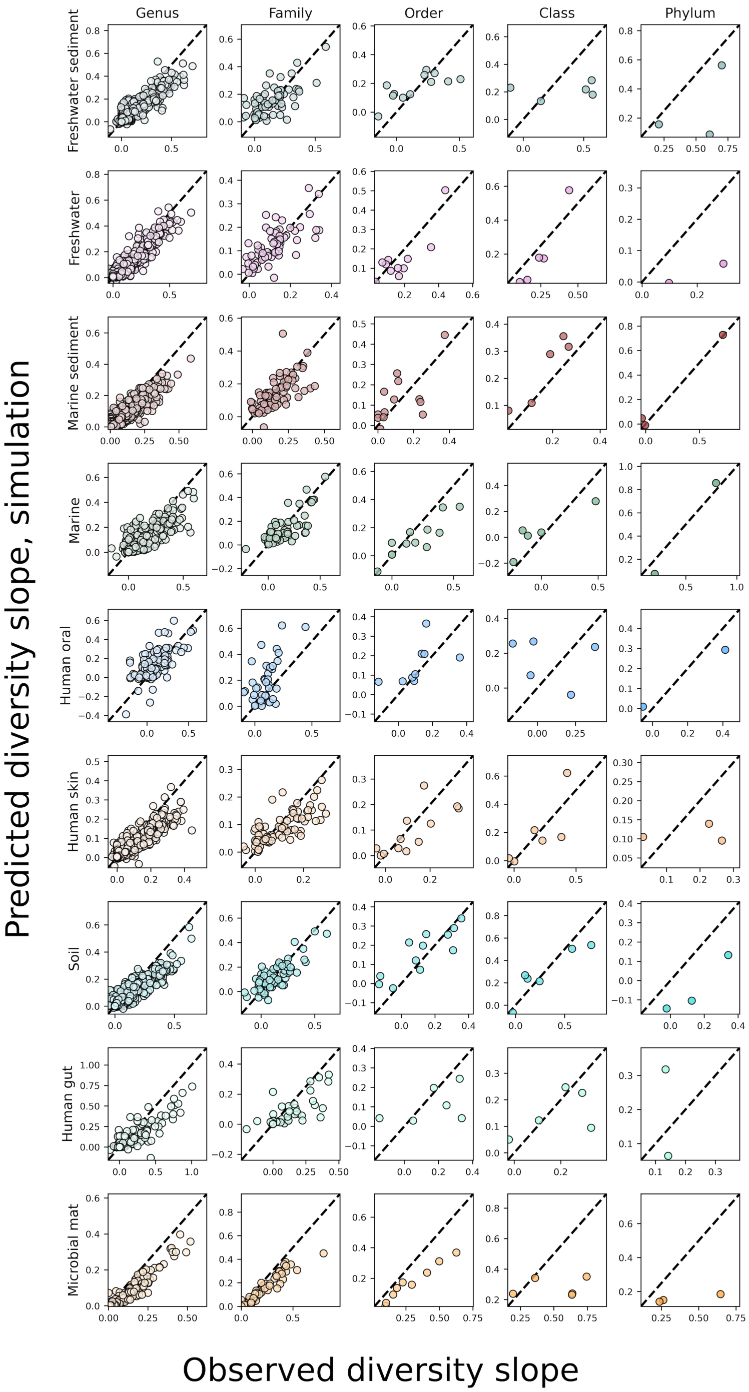


**Figure S31.** The predicted slopes of fine vs. coarse-grained diversity from the sampling form of the gamma distribution with correlations between OTUs under taxonomic coarse-graining.

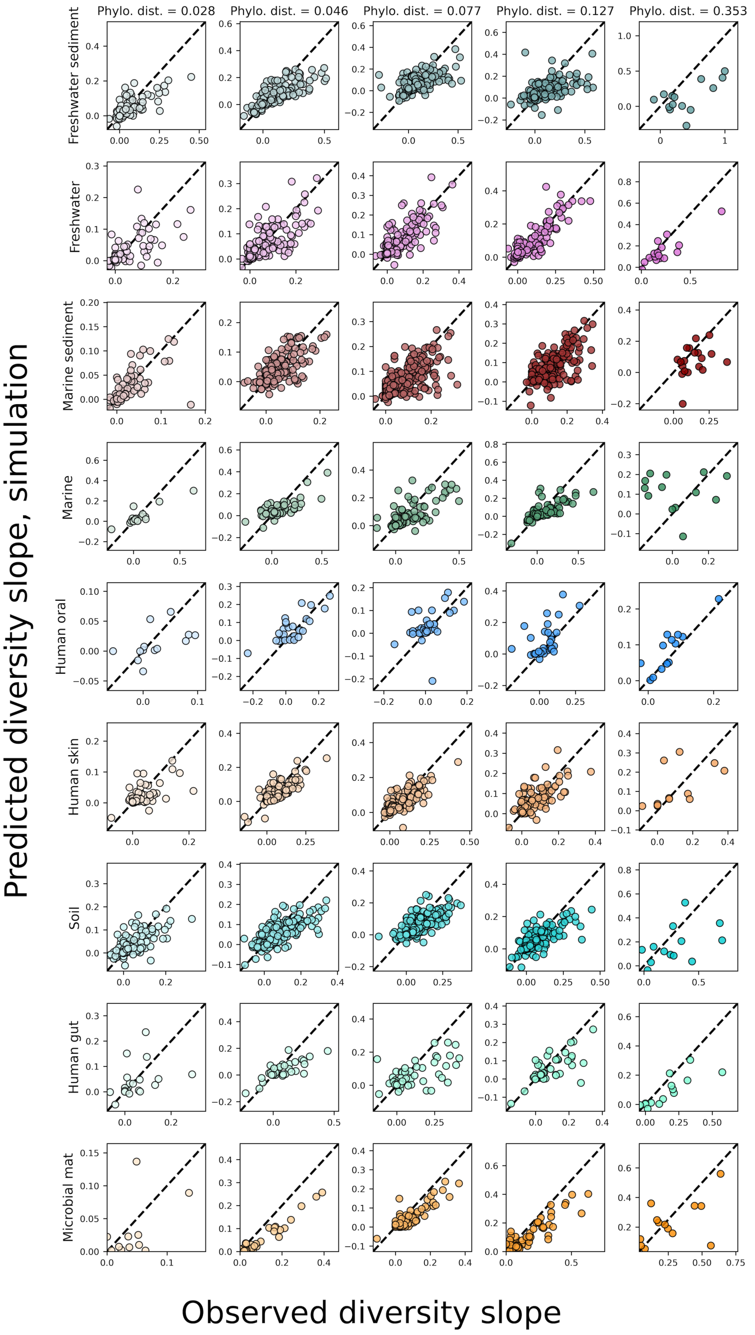


**Figure S32.** The predicted slopes of fine vs. coarse-grained diversity from the sampling form of the gamma distribution with correlations between OTUs under phylogenetic coarse-graining.

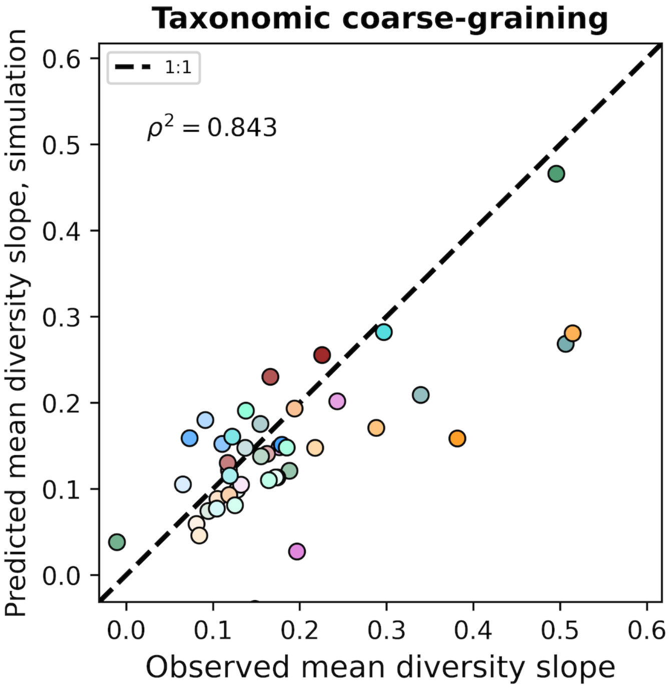


**Figure S33. Gamma distribution simulations with correlations capture observed diversity slopes.** An equivalent set of analyses as depicted in Fig. 6 for taxonomic coarse-graining.

### Tables

| Environment | Total # OTUs | Mean # OTUs | Mean # reads |
|---|---|---|---|
| Marine | 9,090 | 690.80 | 95,129.43 |
| Marine sediment | 16,110 | 1,393.67 | 40,440.38 |
| Human gut | 6,175 | 599.09 | 32,894.50 |
| Human oral | 4,716 | 537.45 | 44,271.22 |
| Human skin | 17,955 | 1,293.45 | 36,344.13 |
| Freshwater sediment | 12,231 | 1,080.95 | 18,979.56 |
| Microbial mat | 5,087 | 200.24 | 8,659.36 |
| Freshwater | 12,052 | 822.37 | 33,646.53 |
| Soil | 20,298 | 1,814.76 | 36,268.93 |

**Table S1.** Summary statistics for the 100 sites randomly selected for each environment. These statistics reflect the data used for taxonomic coarse-graining, as OTUs lacking taxonomic labels were excluded from taxonomic coarse-graining analyses.

| Environment | Total # OTUs | Mean # OTUs | Mean # reads |
|---|---|---|---|
| Marine | 18,173 | 1,356.37 | 168,520.66 |
| Marine sediment | 41,304 | 4,167.25 | 106,166.92 |
| Human gut | 10,190 | 862.73 | 44,031.12 |
| Human oral | 7,062 | 614.97 | 46,104.84 |
| Human skin | 29,448 | 1,817.12 | 48,285.68 |
| Freshwater sediment | 33,193 | 3,569.59 | 65,582.56 |
| Microbial mat | 11,869 | 431.42 | 23,216.02 |
| Freshwater | 26,645 | 1,775.89 | 74,298.17 |
| Soil | 45,273 | 4,730.74 | 106,578.45 |

**Table S2.** Summary statistics for the 100 sites randomly selected for each environment. These statistics reflect the data used for phylogenetic coarse-graining as all OTUs could be used.
